# Histone-organized chromatin in bacteria

**DOI:** 10.1101/2023.01.26.525422

**Authors:** Antoine Hocher, Shawn P. Laursen, Paul Radford, Jess Tyson, Carey Lambert, Kathryn M Stevens, Mathieu Picardeau, R. Elizabeth Sockett, Karolin Luger, Tobias Warnecke

**Author notes:** joint first authors (Antoine Hocher, Shawn P. Laursen).

## Abstract

Histones are the principal constituents of chromatin in eukaryotes and most archaea, while bacteria generally rely on an orthogonal set of proteins to organize their chromosomes. However, several bacterial genomes encode proteins with putative histone fold domains. Whether these proteins are structurally and functionally equivalent to archaeal and eukaryotic histones is unknown. Here, we demonstrate that histones are essential and are major components of chromatin in the bacteria *Bdellovibrio bacteriovorus* and *Leptospira interrogans*. Patterns of sequence evolution suggest important roles in several additional bacterial clades. Structural analysis of the *B. bacteriovorus* histone (Bd0055) dimer shows that histone fold topology is conserved between bacteria, archaea, and eukaryotes. Yet, unexpectedly, Bd0055 binds DNA end-on and forms a sheath of tightly packed histone dimers to encase straight DNA. This binding mode is in stark contrast to archaeal, eukaryotic, and viral histones, which invariably bend and wrap DNA around their outer surface. Our results demonstrate that histones are integral chromatin components across the tree of life and highlight organizational innovation in the domain Bacteria.

## INTRODUCTION

The genomes of all eukaryotes are organized by nucleosomes, which are composed of four core histones that assemble into an octamer to wrap DNA in two tight superhelical turns (Luger *et al*. 1997). This arrangement severely restricts access to the genome. Consequently, eukaryotes rely on a complex system of access control to coordinate DNA-templated processes such as transcription, replication, and DNA repair (Kornberg and Lorch 2020).

Histones are among the most conserved and abundant proteins across eukaryotes (Postberg *et al*. 2010; Patwal *et al*. 2021; Soo and Warnecke 2021). The ‘histone fold’ is composed of three alpha helices (connected by two short loops) and dimerizes in a head-to-tail ‘handshake motif’ (Arents *et al*. 1991). Additional structural elements, including divergent N-terminal tails, distinguish the four eukaryotic core histones from each other (Luger and Richmond 1998a; b). Deletion or depletion of individual histones leads to transcriptional dysregulation, cell cycle arrest, and, ultimately, cell death (Han *et al*. 1987; Kim *et al*. 1988; Wyrick *et al*. 1999; Gossett and Lieb 2012).

Minimalist histones consisting only of the histone fold are pervasive in archaea, where they can play a major role in chromatin organization alongside other nucleoid-associated proteins (NAPs) (Henneman *et al*. 2018; Talbert *et al*. 2019; Laursen *et al*. 2021; Hocher *et al*. 2022). In some archaea, histones rank among the most abundant proteins in the cell (Hocher *et al*. 2022) and their presence is required for viability in the model archaeon *Thermococcus kodakarensis* (Čuboňová *et al*. 2012). Like their eukaryotic counterparts, archaeal histone dimers bend DNA around their outer surface by contacting three consecutive minor grooves of DNA. Three independent DNA interaction interfaces are formed by main and side chains of paired L1-L2 loops and the central α1-α1 interface (Luger and Richmond 1998a; Mattiroli *et al*. 2017). Whereas eukaryotic histones are obligate heterodimers, model archaeal histones such as HTkA/B from *T. kodakarensis* and HMfA/B from *Methanothermus fervidus* (henceforth HMf/HTk histones) can form homo-as well as heterodimers (Sandman *et al*. 1994; Stevens *et al*. 2020). These dimers then oligomerize into a histone stack of variable size that wraps DNA to form slinky-like ‘hypernucleosomes’ (Maruyama *et al*. 2013; Mattiroli *et al*. 2017; Bowerman *et al*. 2021).

Bacteria use a variety of small, basic NAPs to organize their DNA. The most widespread NAP is HU, but other abundant, lineage-restricted proteins are commonplace (Luijsterburg *et al*. 2008; Dillon and Dorman 2010). Deleting individual NAPs is often not lethal to bacterial cells. This is true even for NAPs that, like HU in *E. coli*, are major constituents of the nucleoid (Baba *et al*. 2006). Histones are generally considered to be absent from bacteria. However, homology searches have revealed the presence of proteins with putative histone fold domains in a small and eclectic assortment of bacterial genomes (Alva and Lupas 2018). The role of these proteins in organizing bacterial chromatin is unknown. Here, we show that histone proteins are major and essential nucleoid components in the bacterium *Bdellovibrio bacteriovorus*. In vitro, these histones completely coat linear DNA by binding end-on, forming a nucleohistone filament. In all other known histone-DNA complexes, DNA is wrapped around the outside of a helical ramp of histones. As such, this bacterial ‘inside-out’ arrangement presents a fundamental divergence from the established histone binding mode.

## RESULTS

### Histones are phylogenetically entrenched in several bacterial clades

We conducted a systematic homology search for histone fold proteins in bacteria. Across a phylogenetically balanced database of 18,343 bacterial genomes, we found 416 proteins that contain a histone fold domain (Table S1). 1.86% of genomes encode at least one histone fold domain, compared to 92.8% of genomes that encode HU. In agreement with prior work (Alva and Lupas 2018), we identified two major histone size classes, containing either a singlet or a doublet histone fold. Both classes are typically devoid of other recognized domains (Fig. 1a, Table S1). Like their archaeal homologs, most bacterial singlets also lack the long, disordered N-terminal tails that are characteristic of eukaryotic histones (Henneman *et al*. 2018; Stevens and Warnecke 2022). Amino acid conservation across the three domains of life is particularly high in the L2 loop (RKTV). In archaeal and eukaryotic histones, this region contacts DNA. The highly conserved ‘RD clamp’ stabilizes this arrangement. Bacterial singlets are six residues shorter than HMf/HTk histones, on average. This is mostly due to a shorter α2 helix, which is diagnostic of bacterial singlet histones (Fig. 1b, c). Several residues that are normally present in archaeal and eukaryotic histones, including the ‘sprocket arginine’ R19 (Hodges *et al*. 2015), are not found in bacterial histones (Fig. 1c). The N-terminus as a whole exhibits considerable divergence, is more hydrophobic in nature, and includes a conserved serine-lysine motif (S9-K10) that is not found in archaea.

**Figure 1.**
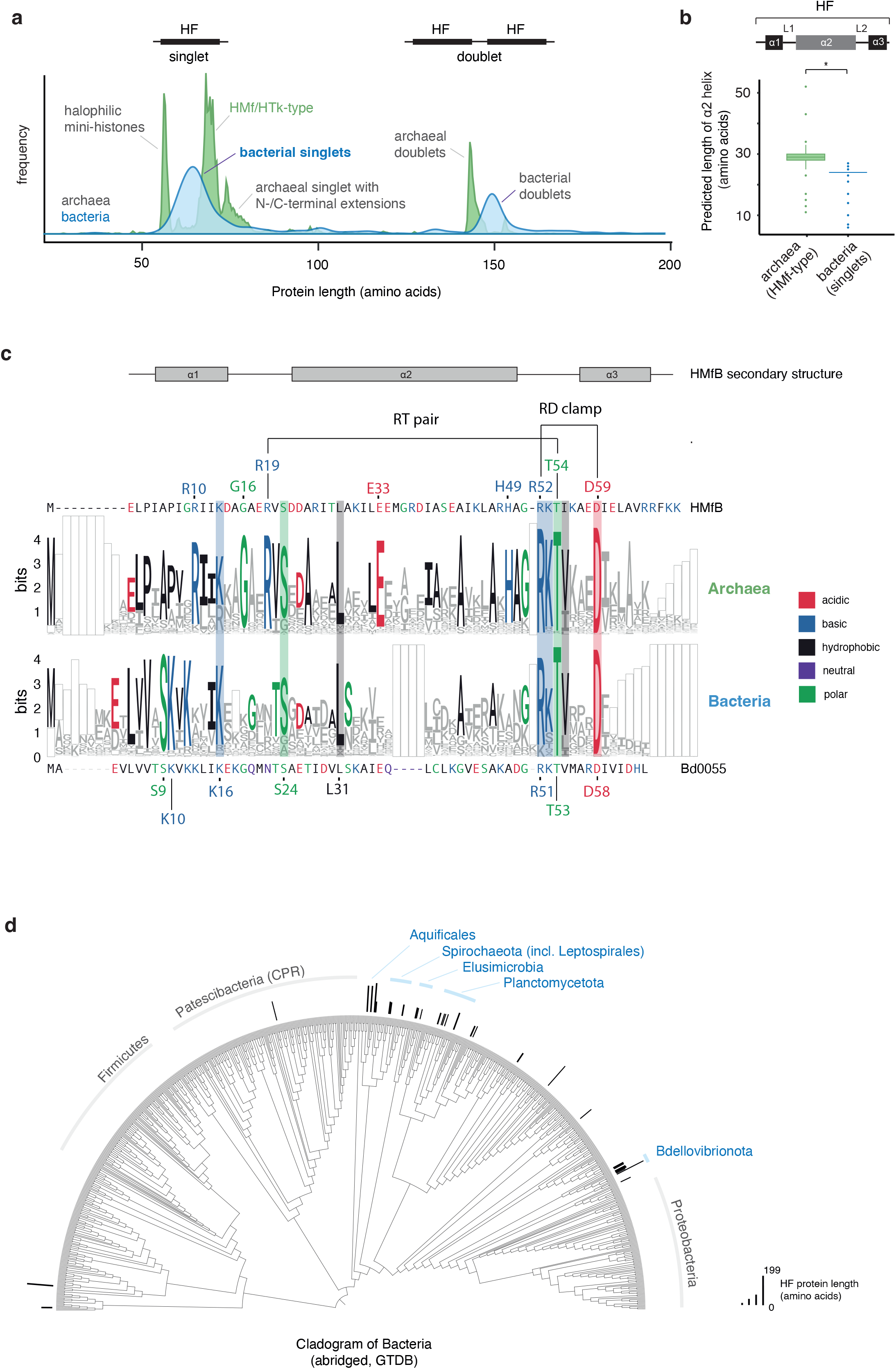
Histone-fold proteins in bacteria. **(a)** Length distribution of proteins (<200 amino acids) encoding predicted histone fold domains in bacteria and archaea. Relative frequencies are shown. **(b)** Length of the α2 helix in bacterial versus HMf/HTk archaeal singlet histones (P<2.2*10-16, Mood test). **(c)** Weblogo representation for bacterial versus archaeal singlet histones. Bacterial and archaeal histones were aligned separately followed by profile-profile alignment to allow comparison across kingdoms. Alignment gaps were coded as a separate character so as to retain their information value and are visualized as empty boxes. Residues that are notably conserved across kingdoms are highlighted by shaded boxes. **(d)** Abridged (see Methods) bacterial species tree illustrating the phyletic distribution of histone fold-containing proteins across the kingdom.

### Bdellovibrio bacteriovorus encodes an abundant histone that localizes to the nucleoid

The phyletic distribution of histones across the bacterial domain is patchy (Fig. 1d, Table S1). In some instances, this might indicate assembly contaminants or recent/transient horizontal gene transfer. However, we identified clades where histone fold proteins are present in several closely related sister lineages. Such phylogenetic persistence is particularly evident in the phylum Bdellovibrionota (Figs. 2a, S1) and the order Leptospirales (Fig. S2).

**Figure 2.**
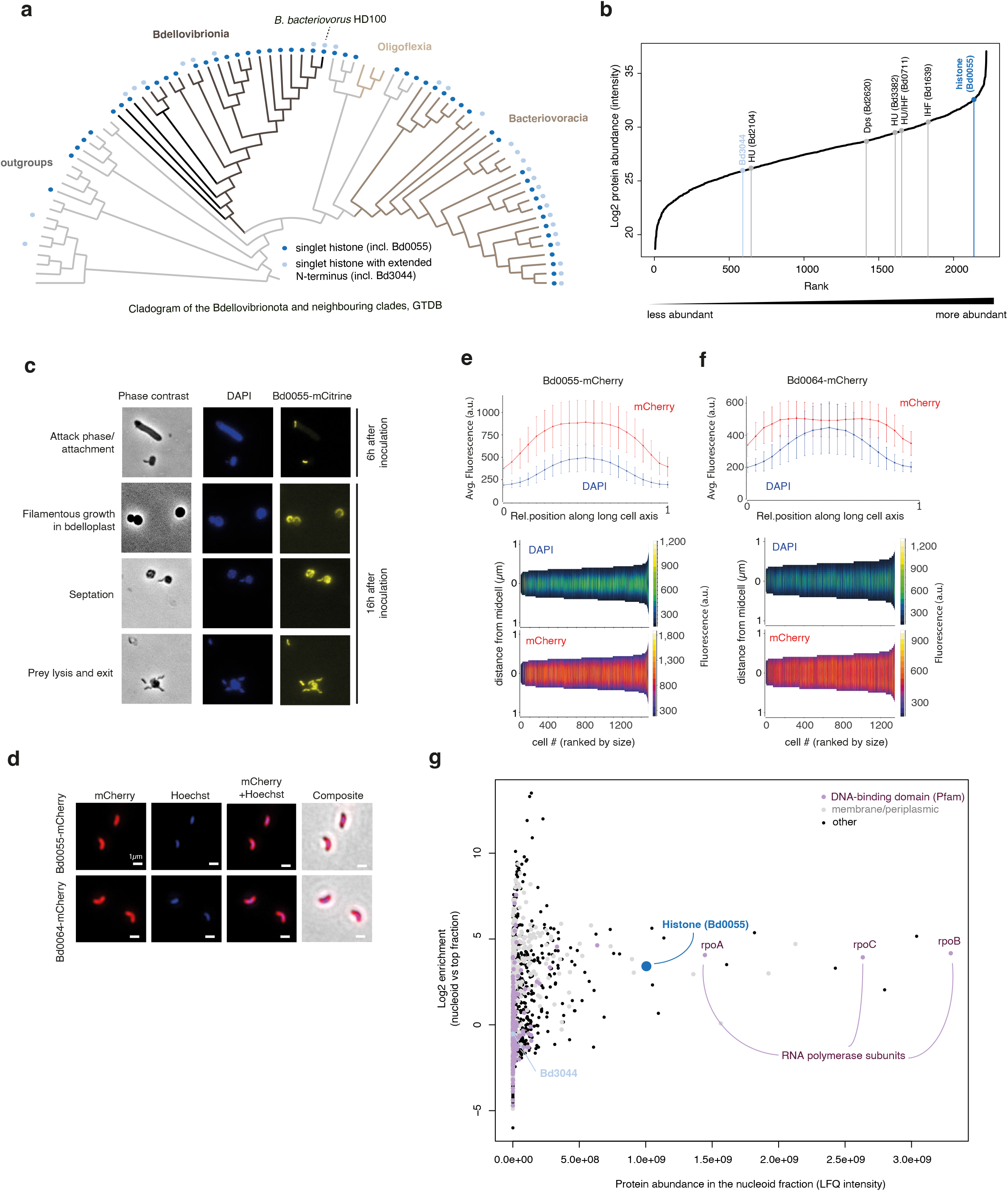
Histones in Bdellovibrio. **(a)** Phyletic distribution of singlet histones across the Bdellovibrionota. **(b)** Ranked protein abundance in *B. bacteriovorus* attack phase cells, based on quantitative label-free proteomics. All quantified proteins are plotted. **(c)** Representative images of different phases of the *B. bacteriovorus* life cycle from strains expressing a Bd0055-mCitrine fusion protein. **(d-f)** Representative images and quantification describing the localization of Bd0055-mCherry and Bd0064-mCherry, a protein with previously established cytosolic localization, in *B. bacteriovorus*. **(g)** High abundance and prominent nucleoid enrichment of Bd0055 in *B. bacteriovorus* attack phase cells (also see Fig. S5). Note that the nucleoid fraction is also enriched for membrane components that have previously been reported to co-sediment with the nucleoid (Portalier and Worcel 1976; Murphy and Zimmerman 1997; Hocher *et al*. 2022).

*Bdellovibrio bacteriovorus* HD100, the model organism of the Bdellovibrionota, is a bacterial predator with a biphasic, bacterially-invasive life cycle (Sockett 2009). Small, motile attack phase cells breach, enter through, and subsequently re-seal the outer membrane of other gram-negative prey bacteria (e.g. *E. coli*). From the periplasm, *B. bacteriovorus* then consumes its prey by secreting proteases and nucleases into the prey cytoplasm, culminating in cycles of replication and coordinated non-binary division to yield new attack phase cells, which are released from the husk of the ravaged prey following induced rupture of its wall and outer membrane (Negus *et al*. 2017; Harding *et al*. 2020).

*B. bacteriovorus* encodes two predicted singlet histones (Table S1): Bd0055 is a bacterial singlet (Fig. 1c) with a detectable homolog in 67% of Bdellovibrionota genomes (Figs. 2a, S1). It is highly conserved at the amino acid level (Fig. S3). Bd3044 is a longer, less well conserved protein that is present in only 37% of Bdellovibrionota genomes (Figs. S1,3). Prior transcriptomic data from across different stages of the *B. bacteriovorus* life cycle indicate high expression of Bd0055, especially during active replication in the host (Fig. S4). We therefore focus on Bd0055 as a candidate global organizer of the nucleoid.

We used quantitative label-free proteomics in attack phase cells to confirm high abundance of Bd0055 at the protein level (Fig. 2b). Of note, protein abundance in attack phase cells is better correlated with RNA abundance in growth phase, consistent with a time lag in protein production where transcript levels during growth phase foreshadow protein levels in attack phase cells (Fig. S4).

To investigate cellular localization of Bd0055 and to monitor expression throughout the *B. bacteriovorus* life cycle, we generated *B. bacteriovorus* HD100 strains in which the native copy of *bd0055* was replaced with a version that was C-terminally tagged with either mCherry or mCitrine. Both tagged strains exhibit no gross morphological defects and carry out predation efficiently. We detect strong fluorescence from tagged Bd0055 throughout the life cycle, including in free-swimming attack phase cells and following invasion of the *E. coli* prey (Fig. 2c). Compared to a previously characterized mCherry-tagged control protein with known cytoplasmic localization (Willis *et al*. 2016; Makowski *et al*. 2019) the fluorescent signal emanating from Bd0055-mCherry is absent from the cell poles, consistent with localization to the nucleoid as captured by Hoechst staining (Fig. 2d-f). In line with the absence of a signal peptide, we find no evidence for secretion into the *E. coli* prey, suggesting that Bd0055 is unlikely to be used for manipulation of prey chromatin.

To obtain orthogonal *in vivo* confirmation that Bd0055 associates with the *B. bacteriovorus* nucleoid, we carried out sucrose gradient-based nucleoid enrichment experiments coupled to quantitative proteomics. We analyzed two fractions from sucrose gradients of lysed attack phase cells: a lower density fraction that is enriched for soluble cytosolic proteins and a nucleoid fraction, enriched for DNA and DNA-binding proteins. Subunits of the RNA polymerase are, as in other bacteria, very abundant (Fig. 2g). Bd0055 is highly represented and enriched 29-fold in the nucleoid fraction. Homologs of classic bacterial NAPs encoded in the *B. bacteriovorus* genome, including HU and Dps, are present but less abundant and less enriched in the nucleoid fraction (Fig. S5).

### Bd0055 folds into a histone fold dimer

We determined the crystal structure of Bd0055 at 1.8 Å resolution (Table S2). Bd0055 forms a crystallographic dimer where one monomer is related to its partner through two-fold symmetry (Fig. 3a). It exhibits the overall topology of a histone fold dimer, as well as a ‘basic ridge’ along its ‘top’ surface that is also found in archaeal and eukaryotic histones, where it is used for DNA binding (Luger *et al*. 1997; Mattiroli *et al*. 2017) (Fig. 3b). The most pronounced structural difference between bacterial and archaeal/eukaryotic histones is the 7 Å shorter α2 helix (1 helical turn shorter compared to archaeal HMfB, Fig. 1c), reducing the overall size of the dimer (Fig. 3a). In addition, the equivalent of the α3 helix in Bd0055 forms only one helical turn and packs against α2 of the dimerization partner. This is unlike the three-turn α3 helix in other histones, which is engaged in tetramerization (Mattiroli *et al*. 2017) by contributing to the four-helix bundle interface, interacting with a conserved histidine in α2 (H49) that is conspicuously absent from all bacterial histones (Fig. 1c). Finally, the surface opposite the basic ridge of the Bd0055 dimer is more acidic than that of HMf/HTk histones (Fig. 3b).

**Figure 3.**
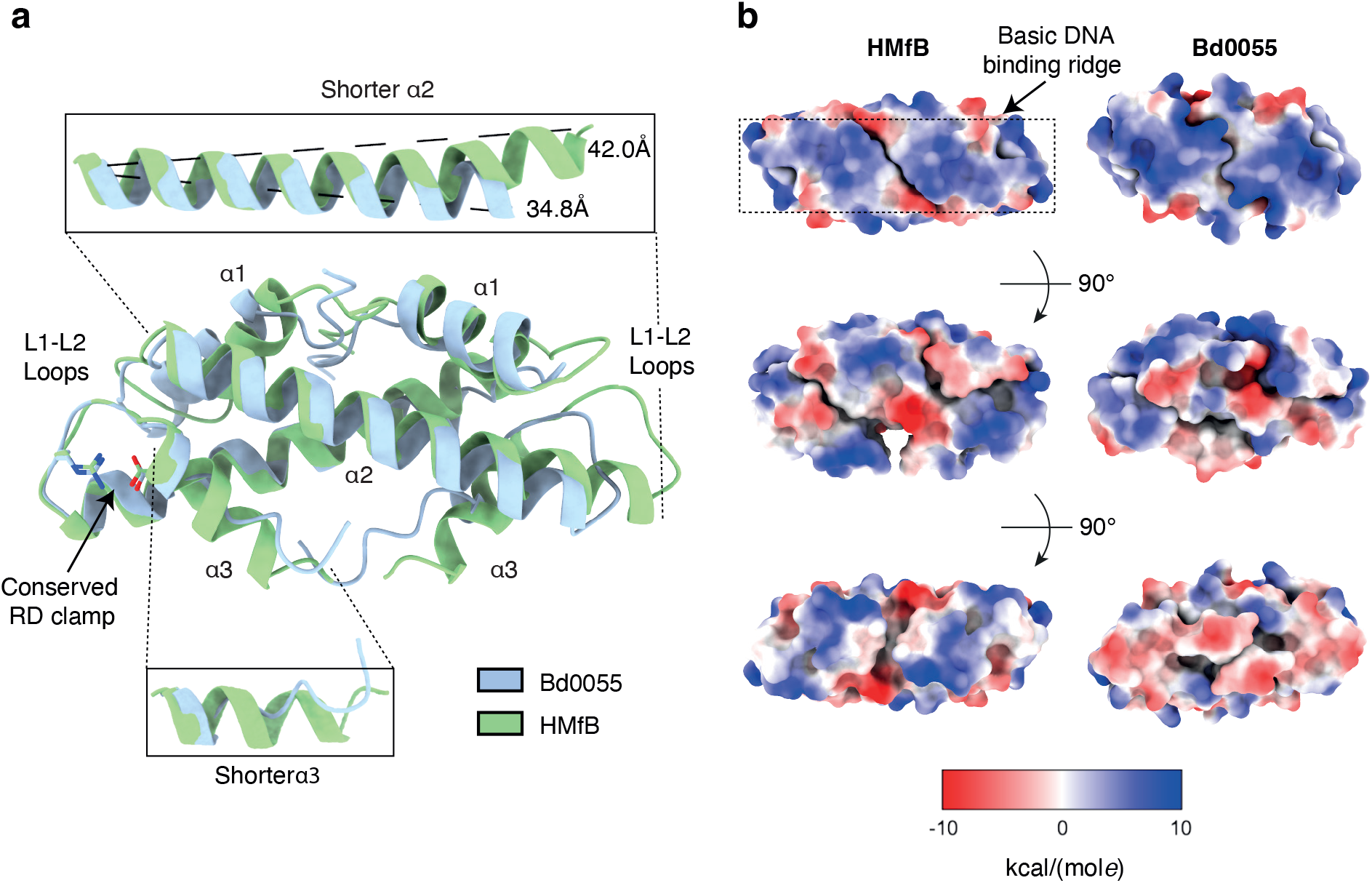
Bd0055 forms a histone fold dimer. **(a)** Crystal structure of Bd0055 superimposed onto archaeal histone HMfB (PDB 1A7W). Bd0055 maintains the overall topology of a histone fold, with shorter *α2* and α3 helices. Bd0055 conserves the signature RD-clamp and L1/L2 loops. **(b)** Columbic surface charge calculation of HMfB and Bd0055, shown in three orientations. Bd0055 maintains the basic ridge of residues important for binding DNA in other histones (top), but has a net acidic charge on its opposite face (bottom).

### Bd0055 binding to DNA differs from that of archaeal histones

We showed that Bd0055 interacts with DNA *in vitro* using fluorescence polarization (FP) (Fig. 4a). Using gel electromobility shift assays, we previously demonstrated that the archaeal histone HTkA shifts 147 bp DNA to a single discrete band (Bowerman *et al*. 2021). HTkA saturates this DNA fragment at a ratio of ~10 to 1 and is unable to produce higher shifts with added protein. In contrast, Bd0055 shifts DNA to several regularly spaced bands indicative of multiple binding events (Fig. 4b). Increasing the protein/DNA ratio of Bd0055 continues to decrease electrophoretic mobility, suggesting a different binding mode (Fig. 4b).

**Figure 4.**
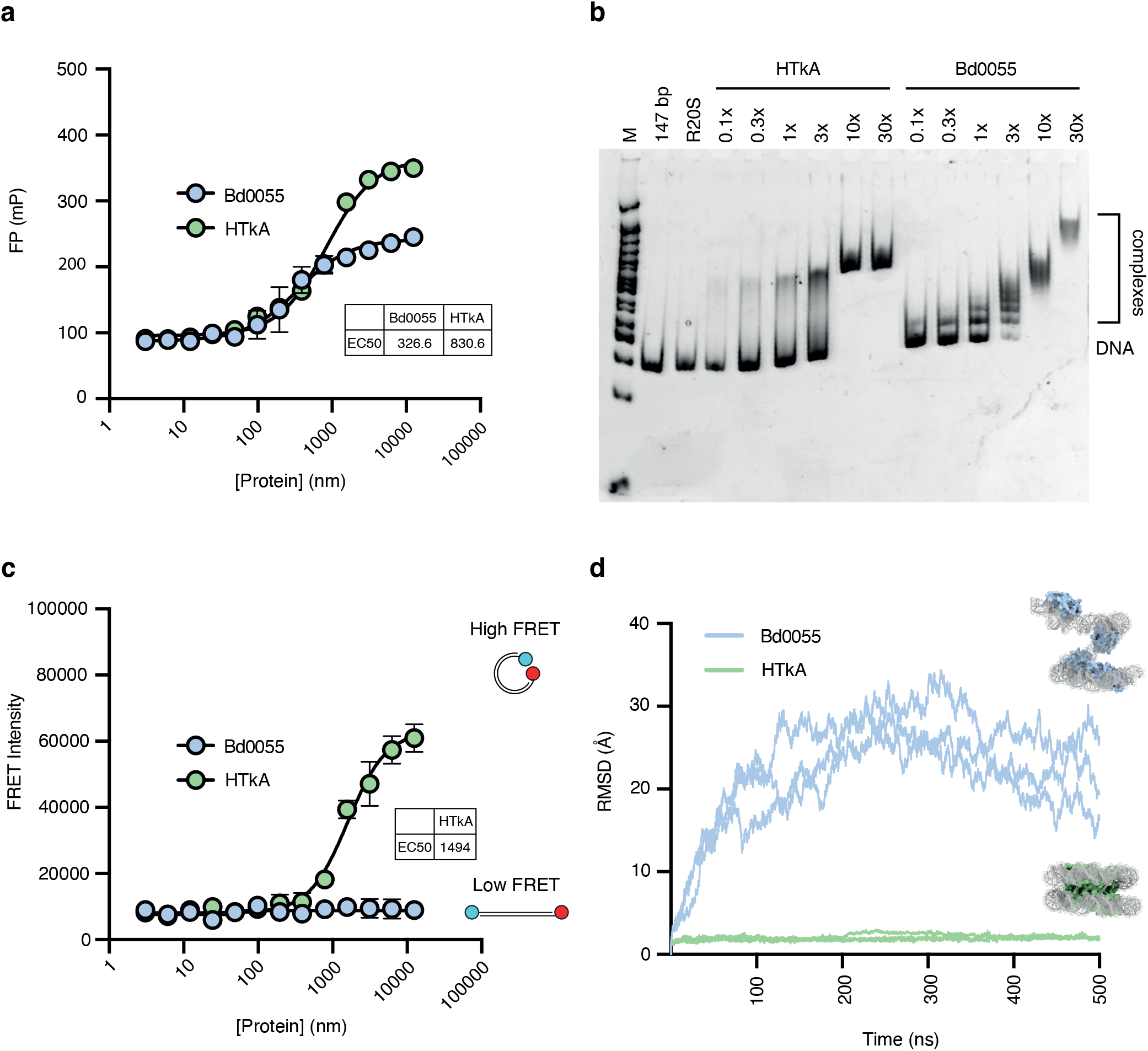
Bd0055 binds DNA differently than HTkA. **(a)** Binding of Bd0055 to an Alexa488-labeled piece of 147 bp DNA (Widom 601 sequence), measured by fluorescence polarization (FP) (n=3). Error bars represent standard deviation of the mean. **(b)** Bd0055 - DNA binding monitored by EMSA (using unlabeled 147 bp DNA). Bd0055 binds with similar affinity as HTkA, but produces a different gel shift pattern. **(c)** Bd0055 was titrated into dual labeled 147 bp DNA (Alexa488 and Alexa647, same DNA as in FP experiment). At concentrations of HTkA high enough to bind DNA, histones wrap the DNA and produce a high FRET signal. Bd0055 does not cause FRET between DNA ends, even with a large excess of protein (n=3). Error bars represent standard deviation of the mean. **(d)** Plot of root mean square deviations (RMSDs) from simulating a hypothetical Bd0055 hypernucleosome modeled onto PDB 5T5K, using four histone dimers and 118 bp of DNA (see Movie S1). After 500 ns of simulation, hypernucleosomes with HMfB remain stable, whereas hypothetical Bd0055 hypernucleosomes fall apart after as little as 10 ns.

To test whether Bd0055 bends DNA into a nucleosome-like geometry, we assembled histones on 147 bp DNA fragments end-labeled with a Förster resonance energy transfer (FRET) donor-acceptor pair. Titration of archaeal HTkA onto this DNA brings the ends into FRET proximity by forming nucleosome-like structures, as observed previously (Mattiroli *et al*. 2017). In contrast, no signal was observed upon titrating even a large excess of Bd0055 (Fig. 4c). By monitoring fluorescence polarization of the same samples, we verified that both proteins bind DNA under these conditions (Fig. 4a). To further investigate this different behavior of Bd0055, we superimposed the Bd0055 histone structure onto an archaeal hypernucleosome consisting of four HMfB dimers and 118 bp of DNA (based on PDB 5T5K) and carried out all-atom molecular dynamics simulations. While archaeal histone dimers remain stably stacked throughout the simulation, the modeled Bd0055 hypernucleosome unfolds within a few nanoseconds (Fig. 4d, Movie S1). Although Bd0055 dimers remain bound to DNA during the simulation, they no longer contact other dimers through protein-protein interactions, suggesting a failure to form stable tetramers on DNA. In archaeal and eukaryotic histones, a tetramer is the smallest unit capable of wrapping DNA.

To understand how Bd0055 binds DNA, we determined the crystal structure of Bd0055 in complex with a 35 bp DNA fragment (Table S2). This structure shows that Bd0055 contacts the DNA with only one L1-L2 binding interface across the minor groove (Fig. 5a). The molecular details of this edge-on interaction are very similar to those observed for other known histones. It employs the L1-L2 motif that is conserved across the domains of life, although the sprocket arginine that reaches into the minor groove in nearly all other histone-DNA complexes is missing (Fig. 1c). Surprisingly, rather than bending the DNA around its positively charged surface, the DNA maintains a straight trajectory. Additional histone dimers bind the next two phosphates through L1-L2 interfaces engaged in identical interactions, but flipped by 180 degrees (Fig. 5b). The filament is stabilized through protein-protein interactions between histone dimers that involve the basic DNA binding ridge and the acidic ‘underside’ of histone dimers 1 and 5, and through exchanging N-terminal tails between dimers 1 and 4 (Fig. 5c). This nucleohistone filament, which requires a histone to DNA ratio of 1 histone dimer per 2.5 bp, reverses known histone convention by completely protecting straight DNA from the solvent. All other known histone-DNA complexes have histones on the inside and wrap DNA around them in a stoichiometry of 1 histone dimer per 30 bp (Fig. 5b).

**Figure 5.**
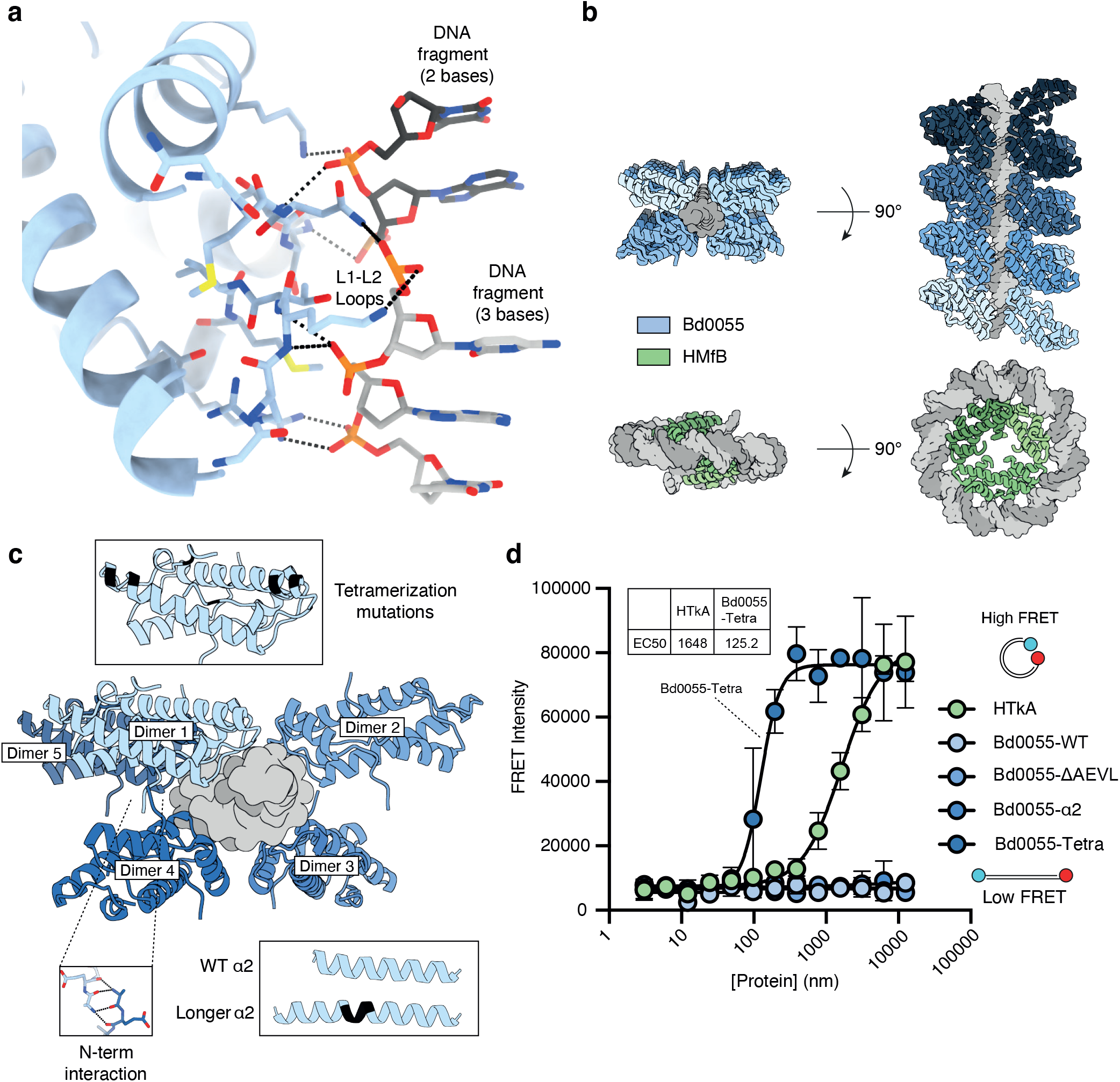
Bd0055 binds DNA end-on and encases straight DNA. **(a)** Crystal structure of Bd0055 in complex with 35 bp DNA, showing interactions between the L1-L2 loop across and the phosphate backbone across the minor groove of DNA. Asymmetric unit is shown. **(b)** Top: Bd0055 dimers encase dsDNA by binding to five phosphates (2-3 on each strand) and interacting with neighboring dimers through electrostatic interactions. Consecutive dimers as they filament on the DNA are shown in decreasing shades of blue. Bottom panel shows how archaeal HMfB wraps DNA around a core of histone dimers that are linked through a four-helix bundle structure (PDB 5T5K). **(c)** Details of histone-histone interactions in the Bd0055 nucleohistone filament. Sites of mutagenesis to switch the binding mode are indicated. **(d)** Mutation of A48H, S45F, I61L (Bd0055-tetra) in Bd0055 enables it to bring DNA ends into FRET proximity, while extending α2 by inserting YAIE (Bd0055-α2) or deleting the N-terminus (Bd0055-ΔAEVL) has no effect, even though all mutants still bind DNA (see Fig. S6). Error bars represent standard deviation of the mean.

### Why does Bd0055 bind DNA edge-on rather than wrapping it around its surface?

Three main structural differences between Bd0055 and other histones could be responsible for this binding mode. First, the N-terminus promotes interactions between dimers on the same strand of DNA, potentially stabilizing the fiber structure. Second, the shorter α2 helix of Bd0055 might force the DNA to bend too severely to wrap around the outside of the shortened dimer. Third, key residues responsible for tetramerization in archaeal histones are absent, most notably a histidine at the equivalent position of A48 (Figs. 1c, 5c).

To test if one of these differences is responsible for the inability of Bd0055 to wrap DNA, we made the compensating mutants *in vitro*, and assayed their ability to wrap 147 bp DNA using FRET. Neither deletion of the four N-terminal amino acids nor insertion of four amino acids into α2 of Bd0055 produce a FRET signal between DNA ends. In contrast, mutations to mimic the archaeal tetramerization domain (A48H, S45F, I61L) enable Bd0055 to bring the ends of the DNA within FRET distance (Fig. 5d). None of these amino acids are implicated in DNA binding in either binding mode and their mutation does not impair the ability of Bd0055 to bind DNA (Fig. S6). The magnitude of the FP signal observed for the tetramerization mutant binding 147 bp DNA closely matches that of HTkA, and the shift to the left indicates the higher affinity of Bd0055 for DNA compared to HTkA (Fig. 5d). These data suggest that the inability of wildtype Bd0055 to form tetramers is responsible for its inability to wrap DNA around its outer perimeter as is observed in archaeal and eukaryotic histones. This is consistent with prior findings from HMfB mutagenesis and analysis of a small set of archaeal histones that also have a non-histidine residue at position 49 and a congruent inability to form stable tetramer interfaces (Marc *et al*. 2002; Stevens *et al*. 2020).

### Bd0055 is essential for predatory and prey-independent growth

We sought to delete *bd0055* using an established silent gene deletion approach to investigate its physiological role (Harding *et al*. 2020; Banks *et al*. 2022). Attempts to delete *bd0055* in the wild type HD100 background resulted in no deletion strains and 133 reversions to wild type. We also attempted to delete *bd0055* in a host-independent strain, HID13 (Lambert *et al*. 2010), to establish whether impaired predation was the reason that no deletions were obtained in HD100. These attempts resulted in no successful deletions and 150 reversions to wild type. This suggests that *bd0055* is essential during both predatory and prey-independent growth.

### Bacterial histones have essential roles outside the Bdellovibrionota

Is *B. bacteriovorus* unique among bacteria in using histones to organize chromatin? We examined publicly available gene expression data and generated additional transcriptomic and proteomic data. At both the transcript and protein level, we find high expression of histone-fold proteins in several distantly related bacteria (Fig. 6a). We also observe notable phylogenetic persistence in clades where gene expression profiles are not available, including in the Planctomycetota and Elusimicrobia (Figs. S7–10).

**Figure 6.**
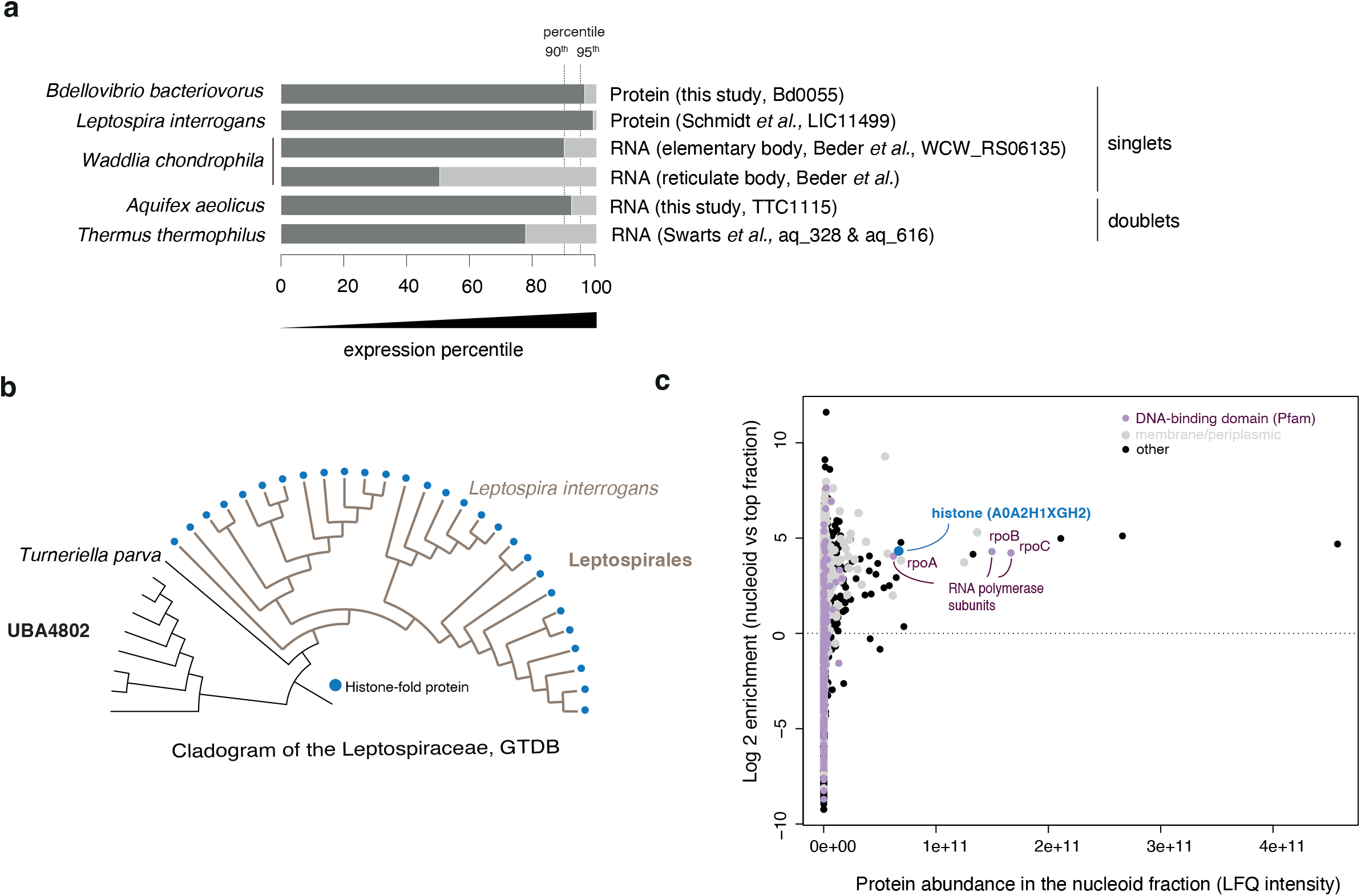
Histones in other bacteria. **(a)** High expression of histone fold proteins at the RNA and/or protein level is evident in bacteria from distant phylogenetic clades. Data sources: *L. interrogans* (Schmidt *et al*. 2011); *Waddlia chondrophila* (Beder and Saluz 2018); *Thermus thermophilus* (Swarts *et al*. 2015). (b) Phyletic distribution of singlet histones across the Leptospirales. (c) High abundance and prominent nucleoid enrichment of the *L. interrogans* histone fold protein (Uniprot ID: A0A2H1XGH2, gene ID: LA_2458).

We became particularly curious about *Leptospira interrogans*, the causative agent of leptospirosis, whose histone (Uniprot ID: A0A2H1XGH2 gene ID: LA_2458) is exceptionally abundant (ranked 4 of 1502 quantified proteins; Figs. 6a, S11). Echoing findings for Bd0055, we find this histone to be highly conserved at the amino acid level with homologs present in all members of the order Leptospirales, including saprophytic and pathogenic species (Figs. 6b, S2,12). Nucleoid enrichment experiments followed by mass spectrometry confirmed high abundance and strong nucleoid enrichment (Fig. 6c). Finally, several attempts to delete the gene encoding this histone from the *L. interrogans* genome failed, indicating that histones are also essential in this species.

## DISCUSSION

Our results demonstrate that histone-based chromatin organization is not exclusive to eukaryotes and archaea. The bacterial histone Bd0055 from *B. bacteriovorus* HD100 is structurally similar to its archaeal and eukaryotic homologs, but binds DNA edge-on and oligomerizes *in vitro* to form a dense nucleohistone fiber that completely encases DNA. This is a striking departure from all histones known to date, which wrap DNA around their exterior to form (hyper)nucleosomes, leaving DNA partially accessible. Our data show that the inability of Bd0055 to form tetramers is responsible for its inability to wrap DNA around its outside. The fact that the tetramerization domain is also absent from all other bacterial singlets (Fig. 1c) further suggests that the inability to wrap might be a general characteristic of bacterial histones.

Filamentation of Bd0055 on DNA is further promoted by complementary charge interactions and hydrogen bonds between individual DNA-bound dimers. Nucleohistone filament formation observed here demands a histone to DNA ratio of 1 histone dimer per ~3 bp of DNA, 10-fold higher than the 1 histone dimer per ~30 bp of DNA observed in canonical nucleosomes and hypernucleosomes. Because crystals were set up at concentrations assuming the much lower stoichiometry, it is unlikely that the nucleohistone filament is an artifact of excessive histones in solution.

The crystal lattice indicates the ability of histones to generate higher order fiber packing through the interaction of histones on adjacent nucleohistone filaments (Movie S2). Although interactions between filaments do not appear to be very strong, it is worth pointing out that one of the three DNA binding regions in the Bd0055 dimer is available for interaction with DNA. Tomograms from *B. bacteriovorus* and *L. interrogans* suggest occasional close packing of DNA fibers *in vivo* (Butan *et al*. 2011; Raddi *et al*. 2012), but whether these are nucleohistone filaments remains to be shown.

The degree of compaction of the *B. bacteriovorus* nucleoid varies throughout its life cycle. Attack phase cells have highly compacted nucleoids that cannot be penetrated even by small fluorescent proteins (Kaljević *et al*. 2021). The nucleoid decondenses shortly after entry into host cells, and a less compact nucleoid state persists until the *B. bacteriovorus* cell is ready to septate. At this time, the newly replicated genomes of each daughter cell again become discretely packaged into highly compacted nucleoids (Borgnia *et al*. 2008). Bd0055 nucleohistone filaments could contribute to nascent genome segregation and the management of extreme compaction during the attack phase. This process could be controlled by changing protein levels, variations in ionic strength, or post-translational modifications.

Histones likely play a role in many other bacteria. Our work shows that histone sequences exist in diverse organisms across the domain Bacteria, are often highly abundant, and are essential in at least one other organism, *L. interrogans*. Like other spirochaetes, *L. interrogans* has a long, slender filamentous shapes, not dissimilar to the shape of *B. bacteriovorus* during predatory growth inside prey, prior to septation. Leptospira histones share key features with Bd0055, such as the shorter α2 helix, the conserved L2 loop, and the absence of the histidine responsible for tetramerization in archaea. More work is needed to discover whether other bacterial histones bind DNA in in the same manner as Bd0055, when and how these sequences were acquired, and what function other bacterial histones that encode doublets or attached domains serve.

## METHODS

### Homology survey of histone fold proteins

PFAM HMM models of known histone fold domains (Histone (PF00125), CBFD_NFYB_HMF (PF00808) and DUF1931 (PF09123)) were searched against a phylogenetically diverse database of 18,343 bacterial genomes (Table S3) using hmmsearch (v 3.1b2) with the –cut_ga option to ensure reproducibility. In addition, a list of eight prokaryotic histone fold seed sequences were obtained from (Alva and Lupas 2018) and used to direct homology search with Jackhmmer (v 3.1b2). Jackhmmer results with p-values <1e-3 were kept. Hits were combined and proteins longer than 200 amino acids discarded. All proteomes in the database were obtained from NCBI (accessed on 20 May 2020). When GenBank proteome files were not available, proteomes were predicted using Prodigal (v2.6.3), with default settings.

For the analysis of α2 length (Fig. 1b), we considered curated sets of histone fold proteins to facilitate the systematic identification of α2. Bacterial singlets were filtered from the bulk of histone fold proteins by removal of DUF1931 HMM hits and further removal of proteins that were longer than 65 amino acids. HMf/HTk-type archaeal histones were pruned from a phylogenetic tree based on the alignment of bacterial and archaeal histone-fold proteins (K. Stevens). Secondary structures were then predicted for each histones using Jpred4 (Drozdetskiy *et al*. 2015) and the length of α2 calculated based on those predictions. The α2 was taken to be the helix that overlapped the peptide region L28-L32 (in HMfB coordinates, see Fig. 1c) and was identified as the longest helix in the protein in 1,439 out of 1,458 cases. Outliers in α2 length (see Fig. 1b) were manually scrutinized and are owing to secondary structure mispredictions, often splitting α2 up into two smaller helices. Removal of these outliers does not affect conclusions.

### Protein alignment and phylogenetic trees

All species trees were obtained from GTDB (https://gtdb.ecogenomic.org/). Trees were rendered by iTol and finalized in Adobe Illustrator.

### Best reciprocal blasts

Best reciprocal blast hits, used to compute amino acid conservation (Figs. S3,12) were obtained using the blast_best_reciprocal_hit function from the R library metablastr (v 0.3), requiring a minimal E-value of 0.001.

### Bacterial culture

*B. bacteriovorus* HD100 was cultured on double-layer YPSC plates to initialize growth from frozen stocks and subsequently grown in liquid calcium/HEPES buffer containing *E. coli* S17-1 as described previously (Lambert and Sockett 2008). The kanamycin-resistant *E. coli* S17-1 (pZMR100) strain was used as prey for the culture of kanamycin-resistant *B. bacteriovorus* strains. Media were supplemented with kanamycin (50 *μ*g/ml) when required. The host-independent *B. bacteriovorus* strain HID13 was grown in YP medium at 30°C as described previously (Lambert *et al*. 2010).

*L. interrogans* was cultured at 30°C in EMJH medium to a density of 10^7^ bacteria/ml. Cells were harvested via centrifugation at 4,000 × g for 20 min and the pellets were washed with Phosphate Buffered Saline (PBS). The resulting pellets were frozen at −80°C until use.

### Whole cell extract preparation

Total proteins were extracted from ~5 mg of −80°C frozen, unfractionated pellet using the iST proteomic kit (see below). Experiments were carried in biological duplicates at the protein extraction step, and each biological replicate was itself technically duplicated at the proteomic step.

### Nucleoid enrichment and protein purification

Nucleoid enrichment was carried out as described in (Ohniwa *et al*. 2011) with the following modifications: Lysozyme concentration in buffer B was doubled for *L. interrogans*; Lysozyme incubation was carried out on ice for 10 min for *B. bacteriovorus* and at room temperature for 10 min for *L. interrogans*. 10 to 60% sucrose gradients were poured manually by 2 mL of 10% increment in Ultra-Clear (14×89 mm) Beckman-Coulter tubes. Gradients were allowed to cool down to 4°C for before the experiment. Lysed cells were deposited on gradients and centrifuged at 10,000 rpm on a SW41-Ti Beckman-Coulter rotor by a Beckman Optima centrifuge pre-cooled at 4°C. Acceleration was set to their minimum value for both the start and the end of the run. Soluble cytosolic proteins settle at low density towards the top of the tube (“top fraction”) whereas the nucleoid, along with membrane proteins (Portalier and Worcel 1976; Murphy and Zimmerman 1997), will settle at a higher density and can be identified as an opaque, viscous band (“nucleoid fraction”). Proteins from both fractions were concentrated and purified following a methanol chloroform treatment as described in (Hocher *et al*. 2022). Samples were then prepared for mass spectrometry using the iST preomics kit, as described below. Experiments were carried out in biological triplicates at the protein extraction step, and each biological replicate was itself technically duplicated at the proteomic step.

### Protein preparation for mass spectroscopy

Proteins from whole cell extract and from different nucleoid enrichment fractions were all processed using the iST 8x Preomics kit. The sonication step was carried out as recommended by the manufacturer on a Bioruptor Plus (high intensity setting). Following heat denaturation at 95°C on an Eppendorf Thermomixer C and sonication, total protein amount was estimated using 205 nm absorbance by Nanodrop (method scope 31). A total of 100 *μ*g for WCE and 37.5 (36.5) *μ*g for *B. bacteriovorus (L. interrogans*) sucrose fractions were used subsequently. Enzymatic digestion (LysC/Trypsin) was carried out on a Eppendorf Thermomixer C for 90 min at 37°C 500 rpm. For each experiment, all samples were processed using the same kit on the same day until storage at −80°C in LC-load buffer.

### Liquid chromatography-tandem mass spectrometry (LC-MS/MS)

Chromatographic separation was performed using an Ultimate 3000 RSLC nano liquid chromatography system (Thermo Scientific) coupled to a Thermo Scientific LTQ Orbitrap Velos (*B. bacteriovorus* nucleoid enrichment samples) or an Orbitrap Q-Exactive mass spectrometer (all other proteomics samples) via an EASY-Spray source.

For sample analysis on the Velos, peptide solutions were injected and loaded onto a trapping column (Acclaim PepMap 100 C18, 100 μm × 2 cm) for desalting and concentration at 8 μL/min in 2% acetonitrile, 0.1% TFA. Peptides were then eluted on-line to an analytical column (EASY-Spray PepMap RSLC C18, 75μm × 50cm) at a flow rate of 250nL/min. Peptides were separated using a 120 minute stepped gradient, 1-22% of buffer B for 90 minutes followed by 22-42% buffer B for another 30 minutes (composition of buffer A – 95/5%: H2O/DMSO + 0.1% FA, buffer B – 75/20/5% MeCN/H2O/DMSO + 0.1% FA) and subsequent column conditioning and equilibration. Eluted peptides were analysed by the mass spectrometer operating in positive polarity using a data-dependent acquisition mode. Ions for fragmentation were determined from an initial MS1 survey scan at 30,000 resolution, followed by CID (Collision Induced Dissociation) of the top 10 most abundant ions. MS1 and MS2 scan AGC targets were set to 1e6 and 3e4 for maximum injection times of 500 ms and 100 ms respectively. A survey scan m/z range of 350 – 1,500 was used, with normalised collision energy set to 35%, charge state screening enabled with +1 charge states rejected and a minimal fragmentation trigger signal threshold of 500 counts.

For sample analysis on the Q-Exactive, chromatographic separation was performed using an Ultimate 3000 RSLC nano liquid chromatography system (Thermo Scientific) coupled to an Orbitrap Q-Exactive mass spectrometer (Thermo Scientific) via an EASY-Spray source. Peptide solutions were injected and loaded onto a trapping column (Acclaim PepMap 100 C18, 100 μm × 2 cm) for desalting and concentration at 8 μL/min in 2% acetonitrile, 0.1% TFA. Peptides were then eluted on-line to an analytical column (EASY-Spray PepMap RSLC C18, 75 μm × 75 cm) at a flow rate of 200 nL/min. See below for further details.

### Specific downstream settings – *B. bacteriovorus* whole cell extract proteomics

Peptides were separated using a 120 minute gradient, 4-25% of buffer B for 90 minutes followed by 25-45% buffer B for another 30 minutes (composition of buffer B – 80% acetonitrile, 0.1% FA) and subsequent column conditioning and equilibration. Eluted peptides were analysed by the mass spectrometer in positive polarity using a data-dependent acquisition mode. Ions for fragmentation were determined from an initial MS1 survey scan at 70,000 resolution, followed by HCD (Higher-energy Collision Induced Dissociation) of the top 12 most abundant ions at 17,500 resolution. MS1 and MS2 scan AGC targets were set to 3e6 and 5e4 for maximum injection times of 50 ms and 50 ms respectively. A survey scan m/z range of 400–1800 was used, normalised collision energy set to 27 and charge exclusion enabled for unassigned and +1 ions. Dynamic exclusion was set to 45 seconds.

### Specific downstream settings – *L. interrogans* proteomics

Peptides were separated using 90-minute gradient, 4-25% of buffer B for 60 minutes followed by 25-45% buffer B for another 30 minutes (composition of buffer B – 80% acetonitrile, 0.1% FA) and subsequent column conditioning and equilibration. Eluted peptides were analysed by the mass spectrometer in positive polarity using a data-dependent acquisition mode. Ions for fragmentation were determined from an initial MS1 survey scan at 70,000 resolution, followed by HCD (Higher-energy Collision Induced Dissociation) of the top 10 most abundant ions at 17,500 resolution. MS1 and MS2 scan AGC targets were set to 3e6 and 5e4 for maximum injection times of 50 ms and 100 ms respectively. A survey scan m/z range of 350–1,800 was used, normalised collision energy set to 27 and charge exclusion enabled for unassigned and +1 ions. Dynamic exclusion was set to 45 seconds.

### Proteomics data processing

Data were processed using the MaxQuant software platform (v1.6.10.43) (Tyanova *et al*. 2016), with database searches carried out by the in-built Andromeda search engine against the GenBank proteome of each organism. A reverse decoy database approach was used at a 1% false discovery rate (FDR) for peptide spectrum matches. Search parameters were as follows: maximum missed cleavages set to 2, fixed modification of cysteine carbamidomethylation and variable modifications of methionine oxidation, protein N-terminal acetylation, asparagine deamidation and cyclisation of glutamine to pyro-glutamate. Label-free quantification was enabled with an LFQ minimum ratio count of 1. The ‘match between runs’ function was used with match and alignment time limits of 0.7 and 20 minutes respectively.

### Nucleoid enrichment analysis

Nucleoid enrichment analysis was carried out as in (Hocher *et al*. 2022), comparing soluble and nucleoid enriched fraction using the R package DEP (v1.8.0). Globally, we observe robust enrichment of proteins with predicted membrane localization, providing validation for the approach (Fig. S13).

### RNA extraction and sequencing

Total RNA was extracted from exponentially growing host-independent *B. bacteriovorus* HID13, and from −80°C frozen *Aquifex aeolicus* pellets (purchased from the Archaeenzentrum Regensburg, Germany) using the RNEasy kit (Qiagen), including DNase I treatment. RNA quality was assessed using an Agilent 2100 Bioanalyser.

For RNA extraction from *B. bacteriovorus* HID13, cells were diluted in 10 mL of fresh YP medium at an OD of 0.1 and grown until reaching a density of 0.6 (20 hours at 30°C with shaking). For each replicate, 2 mL of culture were spun at maximum speed, snap frozen and kept at −80°C until RNA extraction.

For the *B. bacteriovorus* samples (all RIN scores > 9.5), ribosomal RNA was depleted using the NEBNext rRNA Depletion Kit (Bacteria), and NGS libraries made using the NEBNext Ultra II Directional RNALibrary Prep Kit for Illumina according to manufacturer’s instructions. Paired-end 55 bp reads were generated on a NextSeq 2000 with dual 8 bp indexing. For the *A. aeolicus* samples (all RIN scores > 5), rRNA was depleted using Illumina Ribozero Kit (Bacteria), and NGS libraries made using the TruSeq Stranded Total RNA LT Kit according to manufacturer’s instructions. Single End 50 bp reads were generated on a MiSeq with single 6 bp indexing.

### RNA-seq analysis

Untrimmed reads were mapped using Bowtie2 (Langmead and Salzberg 2012) for *A. aeolicus* (single end reads) and using BWA for *B. bacteriovorus* (paired-end). Read counts of all genes were estimated using the python package HTSeq (Anders *et al*. 2015).

### Fluorescence microscopy

The fluorophore mCherry/mCitrine was fused to the C-terminus of Bd0055 by PCR amplification of the gene without its stop codon and amplification of the fluorophore gene, followed by Gibson cloning using the Geneart assembly kit (Invitrogen) according to instructions into the mobilisable broad host range vector pK18*mobsacB* and this was conjugated into *B. bacteriovorus* HD100 as described previously (Lambert *et al*. 2003). PCR amplification was carried out with phusion polymerase (New England Biolabs) according to instructions, using primers: Bd0055mC_mCit_F 5’ cgttgtaaaacgacggccagtgccaACACTCATCATACCCGCC 3’; Bd0055mC_Link_R 5’ TGCTCACCATACCACCGCTGCCACCGCCGCCGCTACCGCCAAGGTGGTCG 3’; 0055mCherry_Link_F 5’ CGACCACCTTGGCGGTAGCGGCGGCGGTGGCAGCGGTGGTATGGTGAGCA 3’; 0055mCherry_R 5’ ggaaacagctatgaccatgattacgTTACTTGTACAGCTCGTCCATG 3’.

Cells were imaged on a Nikon Ti-E epifluorescence microscope equipped with a Apo x100 Ph3 oil objective lens (NA: 1.45) and images were acquired on an Andor Neo sCMOS camera using Nikon NIS software. The following filters were used for fluorescence images: mCherry (excitation: 555 nm, emission: 620/60 nm), DAPI (Hoechst 33342): (excitation: 395 nm, emission: 435-485 nm). DNA was stained by the addition of Hoechst 33342 at a final concentration of 5 *μ*g/ml. Images were analysed with FIJI software with the MicrobeJ plugin (Ducret *et al*. 2016). mCitrine expressing cells were imaged on a Leica DMRB with phase contrast and DIC for transmitted light illumination. To image mCitrine during predation, cells were fixed (1% paraformaldehyde, 5 min, quenched using 150 mM glycine) and DNA stained by DAPI in 1x PSB (5min of staining removal by centrifugation).

### Protein purification from *E,coli*

The Bd0055 (or mutant) ORF was codon optimized for expression in *E. coli* and synthesized into a dsDNA gBlock (IDT) with 35 bp and 22 bp overhangs on the 5’ and 3’ sides, respectively. The gBlock was cloned into a *lac*-inducible, ampicillin-resistant pET expression vector through restriction-free cloning. The protein was expressed in BD *E. coli* cells from overnight cultures using 0.4 mM IPTG induction after cells reached an OD of ~1.0. Cells were harvested after 2 hours and centrifuged at 6,000 rpm for 20 min at 4°C. Media was decanted and the cells were flash frozen in liquid nitrogen and stored at −80°C. Cells were thawed on ice for 30 min and resuspended in lysis buffer (50 mM Tris pH 7.5, 5 mM EDTA, 0.1% Triton X-100, 5 mM BME, 1 mM AEBSF, and 1 Pierce Complete protease inhibitor tablet per 50 ml). The cells were then lysed by sonication (3 rounds of 1s on/off for 1min) and centrifuged at 16,000 rpm for 20 min at 4°C. The lysate was filtered through a 0.45 um filter and run over a 5 ml SP HP column (Cytiva) on an AKTA Pure FPLC using a linear gradient starting at buffer A (0 M NaCl, 50 mM Tris-HCl pH 7.5, 1 mM TCEP, 1 mM AEBSF) to 1 M NaCl (1 M NaCl, 50 mM Tris-HCl pH 7.5, 1 mM TCEP, 1 mM AEBSF) over 40 column volumes. Protein typically eluted around 300 mM NaCl. Fractions were sampled uniformly and analyzed by SDS-PAGE, as Bd0055 has very little absorption at 280 nm. Fractions containing Bd0055 were pooled and diluted in buffer A to ~100 mM NaCl before being applied to a 5ml Heparin HP column (Cytiva). Protein was eluted with a linear gradient from buffer A to buffer B over 40 CV, with the protein typically eluting at 450 mM NaCl. Fractions were pooled and concentrated to a volume of 1 ml before being loaded onto a 120 ml S75 (Cytiva) ran in buffer B containing 10% glycerol, with the protein having a typical retention volume of 74-82 ml. Fractions were pooled, concentrated, aliquoted, flash frozen in liquid nitrogen, and stored at −80°C.

### Apo protein crystallization

Recombinant Bd0055 in storage buffer (1 M NaCl, 1 mM EDTA, 50 mM HEPES pH 7.5, 10% glycerol) was crystallized through hanging drop diffusion in a mother liquor (3.2 M (NH_4_)_2_SO_4_, 100 mM BICINE pH 9) by gently mixing 1 *μ*l of protein to 1 *μ*l of mother liquor on a coverslip and sealing it onto a pre-greased well of a 24 well crystal tray containing 1 ml of mother liquor. Crystals formed within 24 hrs and matured within 3 days. Crystals were looped and frozen in liquid nitrogen. X-ray diffraction data were collected on a Rigaku XtaLAB MM003 and indexed in DIALS. Molecular replacement was done using Phenix and structures were refined in Coot. Short, idealized alpha-helices were used as search models, most of the remaining structure was built automatically using AutoBuild. Final residues were placed by hand in Coot.

### DNA binding (FP)

50 *μ*M protein was titrated using an Opentrons OT-2 liquid handling robot into 10 nM Alexa488 end-labeled 147 bp DNA by serial dilution down to a protein concentration less than the DNA probe in reaction buffer (10 mM NaCl, 1 mM EDTA, 50 mM HEPES pH 6). After incubation at RT for > 15 min (overnight incubation did not change overall results), Fluorescence polarization data were collected using a BMG Labtech CLARIOstar. Data were analyzed in Prism and fit to an [Inhibitor] vs. response – four parameter nonlinear regression curve.

### DNA binding (FRET)

FRET experiments were conducted simultaneously with the FP experiments, as the 147 bp DNA used was labeled on the opposite end of the Alexa488 with an Alexa647 fluorophore. Data were collected on a BMG Labtech CLARIOstar through excitation of 488 nm light while recording the emission at 647 nm. Data were analyzed in Prism and fit to an [Inhibitor] vs. response – four parameter nonlinear regression curve.

### Electrophoretic mobility shift assay (EMSA)

1 *μ*M 147 bp DNA was mixed with varying concentrations of protein in reaction buffer (10 mM NaCl, 1 mM EDTA, 50 mM HEPES pH 7.5). Reactions were mixed 1:1 with 80% glycerol, incubated at RT for > 15 min, and ran on 10% native PAGE (0.2x TBE running buffer, 150 V, 90 min). Gels stained with ethidium bromide and visualized on a Typhoon, using the appropriate laser and filter.

### Molecular dynamics simulations

The HMfB hypernucleosome structure was built in ChimeraX and Chimera, by extending PDB 5T5K to include four HMfB dimers and 118 bp of dsDNA. The Bd0055 hypernucleosome structure was built by docking the Bd0055 apo structure into the HMfB hypernucleosome. All-atom molecular dynamics simulations using explicit solvent were carried out using AMBER18 using the ff14SB, bsc1, and tip3p forcefields (for protein, DNA, and water respectively). Structures were protonated and hydrogen mass repartitioned (as implemented through parmed in AMBER). Structures were placed in cubic boxes surrounding the structures by at least 25 Å, charge neutralized using potassium ions, and hydrated with water molecules. The structures were energy minimized in two, 5,000 step cycles: the first restraining the protein and DNA molecules to allow the solvent to relax, and the second to allow the whole system to relax. Minimized structures were then heated to 300 K and slowly brought to 1.01325 atm. These systems were then simulated for 500 ns in 4 fs steps. Simulations were carried out on NVIDIA GPUs (RTX6000s or A100s) using CU Boulder’s Blanca Condo cluster. RMSD analysis was carried out using cpptraj through AMBER18.

### DNA/protein crystallization

DNA fragments ranging between 35 bp and 45 bp in length, with randomized sequence and ~50% GC content, were screened for their ability to form discrete complexes with Bd0055, using EMSA. The 35 bp DNA (sequence: 5’TCTTGCACTAAGAGCTACTGGAGTGCGTCAGATGT3’) was selected as it formed a discrete DNA shift at roughly 4:1 protein to DNA (it shifted to higher smears upon addition of more protein). 75 *μ*M DNA and 1200 *μ*M protein (16:1) were mixed in crystallization buffer (10 mM NaCl, 0.1 mM EDTA, 50 mM HEPES pH 7.5) and dialyzed at RT against crystallization buffer. Hanging drop crystals were set in 4 *μ*l drops at 1:1 complex to mother liquor (15% PEG 550 MME, 50 mM HEPES pH 8.0) and incubated at 20°C. Crystals formed overnight and matured within a week. Large cubic crystals were looped, cryoprotected by washing in 30% glycerol, and flash frozen in liquid nitrogen. X-ray diffraction data were collected at the ALS synchrotron (12,397.9 eV, 225 mm detector, ΔΦ = 0.25) and indexed in DIALS. Molecular replacement was done using Phenix and structures were refined in Coot. The apo Bd0055 structure was used as a search model along with 2 ssDNA fragments (2 and 3 bp).

### Gene deletion

Attempts to generate a markerless deletion mutant of the *bd0055* open reading frame in *B. bacteriovorus* HD100 were by PCR amplification of the flanking 1000 bp upstream region with the first 2 codons of the *bd0055* open reading frame and final 3 codons with the downstream 1000 bp flanking region and fusing these by Gibson assembly into the mobilisable broad host range vector pK18*mobsacB*. The primers used were: Bd0055_UP_F 5’ cgttgtaaaacgacggccagtgccaATTGATTGAACACGGCAATC 3’; Bd0055_UP_R 5’ cgaattaaaggtgTGCCATGATATACCTCTCC 3’; Bd0055_DN_F 5’ gtatatcatggcaCACCTTTAATTCGGTCGAACG 3’; Bd0055_DN_R 5’ ggaaacagctatgaccatgattacgTCGCTGGATGTCTCCGGC 3’ This construct was conjugated into *B. bacteriovorus* HD100 as described previously (Lambert *et al*. 2003) and resulting exconjugants screened for kanamycin sensitivity and screened by PCR to test for either gene deletion or revertant to wild-type. Screening primers were: Bd0055KO_S_F 5’ atctggagcttcacttcccg 3’and Bd0055KO_S_R 5’ ggtgatgatccgggctctaa 3’.

For targeted mutagenesis in *L. interrogans*, a kanamycin resistance cassette replacing the coding sequence of LA_2458 and 0.8-0.9 kb sequences homologous to the sequences flanking the target gene was synthesized by GeneArt (Life Technologies, Grand Island, NY, USA) and cloned in an *E. coli* vector which is not able to replicate in *L. interrogans*. Plasmid DNA was then introduced in *L. interrogans* serovar Manilae by electroporation as previously described (Picardeau *et al*. 2001) with a Biorad Gene Pulser Xcell™. Electroporated cells were plated on EMJH agar plates supplemented with 50 μg/ml kanamycin. Plates were incubated for 4 weeks at 30°C in sealed plastic bags or wrapped in foil to avoid desiccation.

## Dataset availability

All datasets are publicly available. Proteomics data have been deposited to PRIDE (PXD039405) and RNA-seq data to GEO under accession GSE220534. All genomes used were publicly available with no usage restriction (Table S3). Structure data have been deposited at the Protein Data Bank (PDB 8FVX for apo Bd0055 and 8FW7 for DNA bound Bd0055).

## Script availability

Analysis scripts were run using R version 3.6.2. Scripts required to reproduce figures are available at https://github.com/hocherantoine/BHF/.

## ACKNOWLEDGMENTS

We thank the LMS Proteomics Facility for generating mass spectrometry data, the LMS Genomics Facility for RNA sequencing, and Annette Erbse at CU Boulder for help with crystallography. PR, JT, CL, and RES are funded by a Wellcome Trust Investigator Award in Science (209437/Z/17/Z). Work by AH, KMS, and TW is funded by a UKRI MRC core grant to TW (MC-A658-5TY40). KL and SL are funded by the Howard Hughes Medical Institute.

## OPEN ACCESS

For the purpose of open access, the author has applied a Creative Commons Attribution (CC BY) licence (where permitted by UKRI, ‘Open Government Licence’ or ‘Creative Commons Attribution No-derivatives (CC BY-ND) licence may be stated instead) to any Author Accepted Manuscript version arising’.

This article is subject to HHMI’s Open Access to Publications policy. HHMI lab heads have previously granted a nonexclusive CC BY 4.0 license to the public and a sublicensable license to HHMI in their research articles. Pursuant to those licenses, the author-accepted manuscript of this article can be made freely available under a CC BY 4.0 license immediately upon publication.

## MATERIALS & CORRESPONDENCE

Correspondence and requests for materials can be addressed to KL or TW.

## CONTRIBUTIONS

AH initiated the project and carried out all bioinformatic analyses, sequencing experiments and proteomics (including upstream bacterial culture and biochemistry). SL performed all *in vitro* work on *E. coli*-expressed Bd0055 (biophysical characterization, crystallography, structure analysis) as well as MD simulations. KMS assisted in histone homolog curation and pan-bacterial analysis of histone proteins. PR built Bd0055-mCherry and -mCitrine fusion strains of *B. bacteriovorus* which were phenotyped microscopically by JT. CL and PR attempted gene deletions of *bd0055* in predatory and prey-independent cultures. MP provided biomass for *L. interrogans* biochemistry and attempted gene deletions of LA_2458. RES, KL and TW supervised the project. AH, SL, KL, and TW led data analysis and interpretation and wrote the manuscript with input from all authors.

## SUPPLEMENTARY TABLES

**Table S1. Bacterial histone-fold proteins identified via homolog search.**

**Tables S2.** X-ray diffraction statistics. Data collection and refinement statistics from Bd0055 apo crystal structures (pdb 8FVX) and Bd0055 bound to 35 bp of dsDNA (pdb 8FW7).

|  | Bd0055 apo | Bd0055 bound to 35 bp dsDNA |
| --- | --- | --- |
| **Data collection** |  |  |
| Space group | P 2 21 21 | I 21 21 21 |
| Cell dimensions |  |  |
| *a*, *b*, *c* (Å) | 26.7722 34.4118 56.484 | 33.081 103.143 111.145 |
| $\alpha$, $\beta$, $\gamma$ (°) | 90 90 90 | 90 90 90 |
| Resolution (Å) | 29.39 - 1.8 (1.865 - 1.8) | 27.79 - 2.0 (2.072 - 2.0) |
| *R*<sub>merge</sub> | 0.05163 (0.1029) | 0.06045 (0.5857) |
| *I* / $\sigma I$ | 26.79 (10.42) | 16.49 (3.86) |
| Completeness (%) | 99.42 (95.25) | 98.37 (97.89) |
| Redundancy | 8.4 (5.2) | 7.1 (7.4) |
| **Refinement** |  |  |
| No. reflections | 5,156 (461) | 13,136 (1,302) |
| *R*<sub>work</sub> / *R*<sub>free</sub> | 0.1889 (0.2095) / 0.2331 (0.2670) | 0.2218 (0.2320) / 0.2536 (0.2508) |
| No. atoms |  |  |
| Macromolecules | 476 | 1051 |
| Ligand | 0 | 0 |
| Solvent | 43 | 52 |
| Protein residues | 63 | 126 |
| *B*-factors |  |  |
| Macromolecules | 11.88 | 46.07 |
| Solvent | 20.65 | 42.45 |
| R.m.s. deviations |  |  |
| Bond lengths (Å) | 0.007 | 0.008 |
| Bond angles (°) | 0.82 | 0.92 |

Values in parentheses are for highest-resolution shell. One crystal was used for each structure.

**Table S3. Bacterial genomes searched for histone fold proteins.**

## SUPPLEMENTARY FILES

**Movie S1.** 500 ns simulations of HMfB and modeled Bd0055 hypernucleosomes.

**Movie S2.** Animation of nucleohistone filament and higher-order lattice formation. In the crystal lattice, histone-bound DNA fibers assemble in a parallel manner with a DNA spacing of ~60 Å to the nearest strand. The protein-protein contacts stabilizing the lattice engage the second L1-L2 interface of a given Bd0055 dimer. Arginine 57 is sandwiched between E36 and D29 of a histone dimer binding to another DNA, and C-terminal tails cross over between the two dimers (Fig. 5c). Additionally, the amino terminal NH3 hydrogen bonds with D49 in the L2 loop of the neighboring dimer.

---

**Figure S1.**
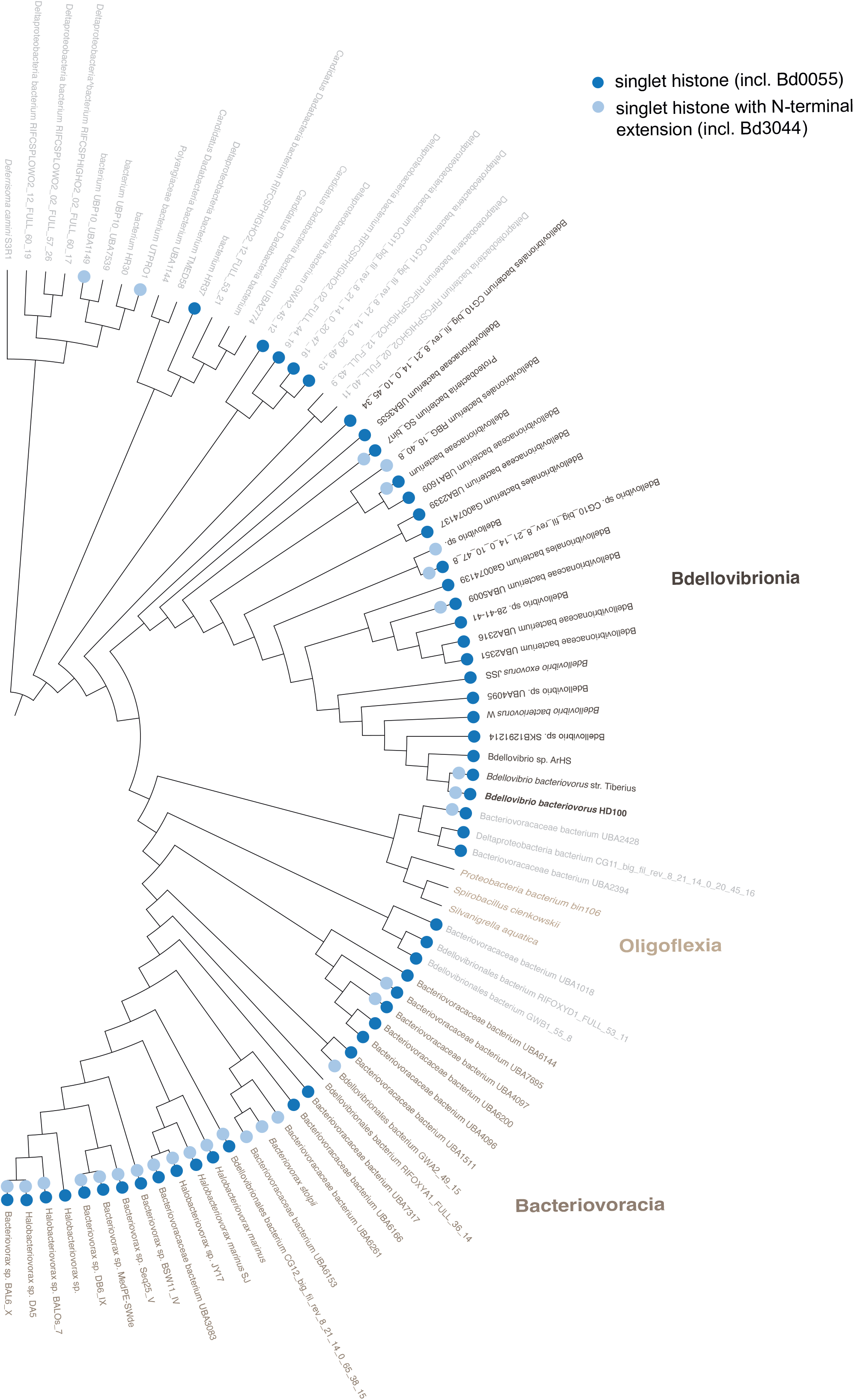
Cladogram of the Bdellovibrionota, indicating the presence of histone fold proteins throughout the clade. This is a fully annotated version of Fig. 2a. The cladogram is based on the publicly available GTDB tree (v 95.0) and represents the subset of genome assemblies that were used in this study and are also present in GTDB.

**Figure S2.**
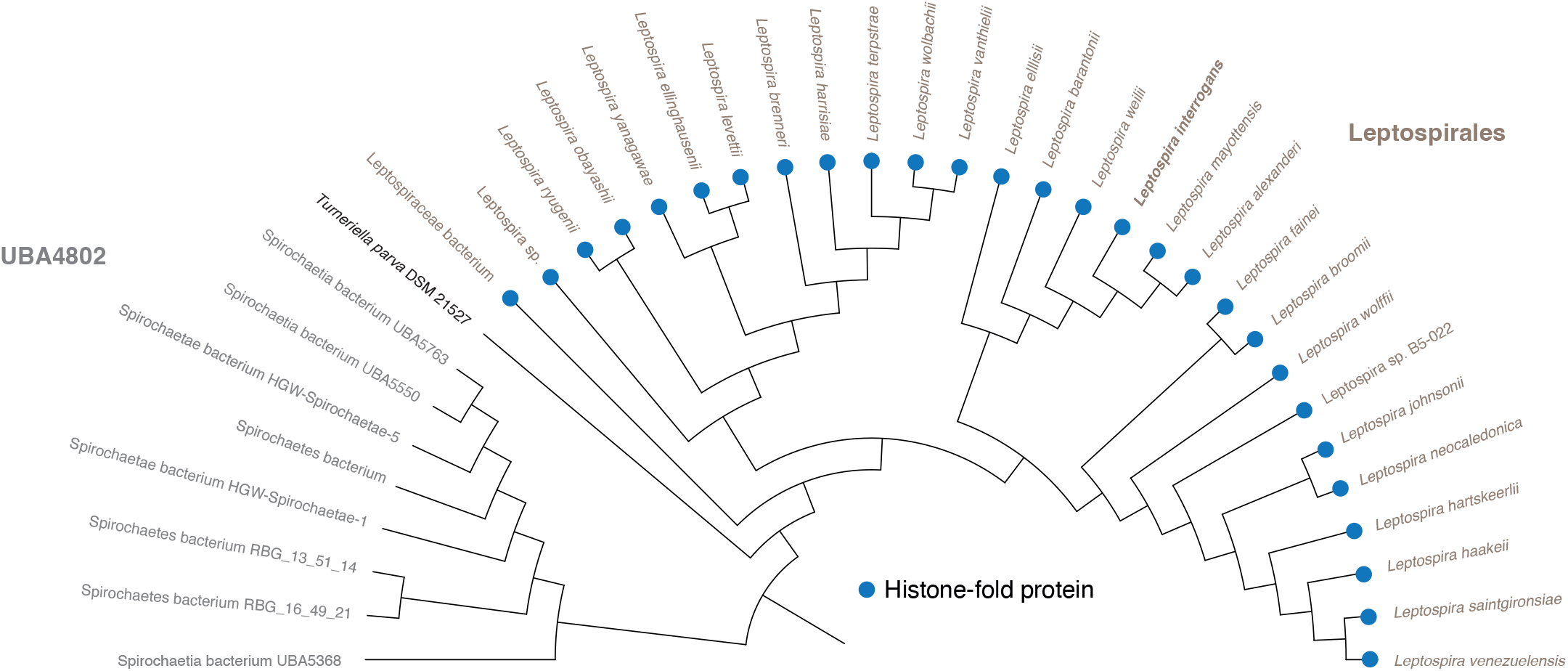
Cladogram of the class Leptospirales and its UBA4802 sister clade, indicating the presence of histone fold proteins throughout the clade. The cladogram is based on the publicly available GTDB tree (v 95.0) and represents the subset of genome assemblies that were used in this study and are also present in GTDB.

**Figure S3.**
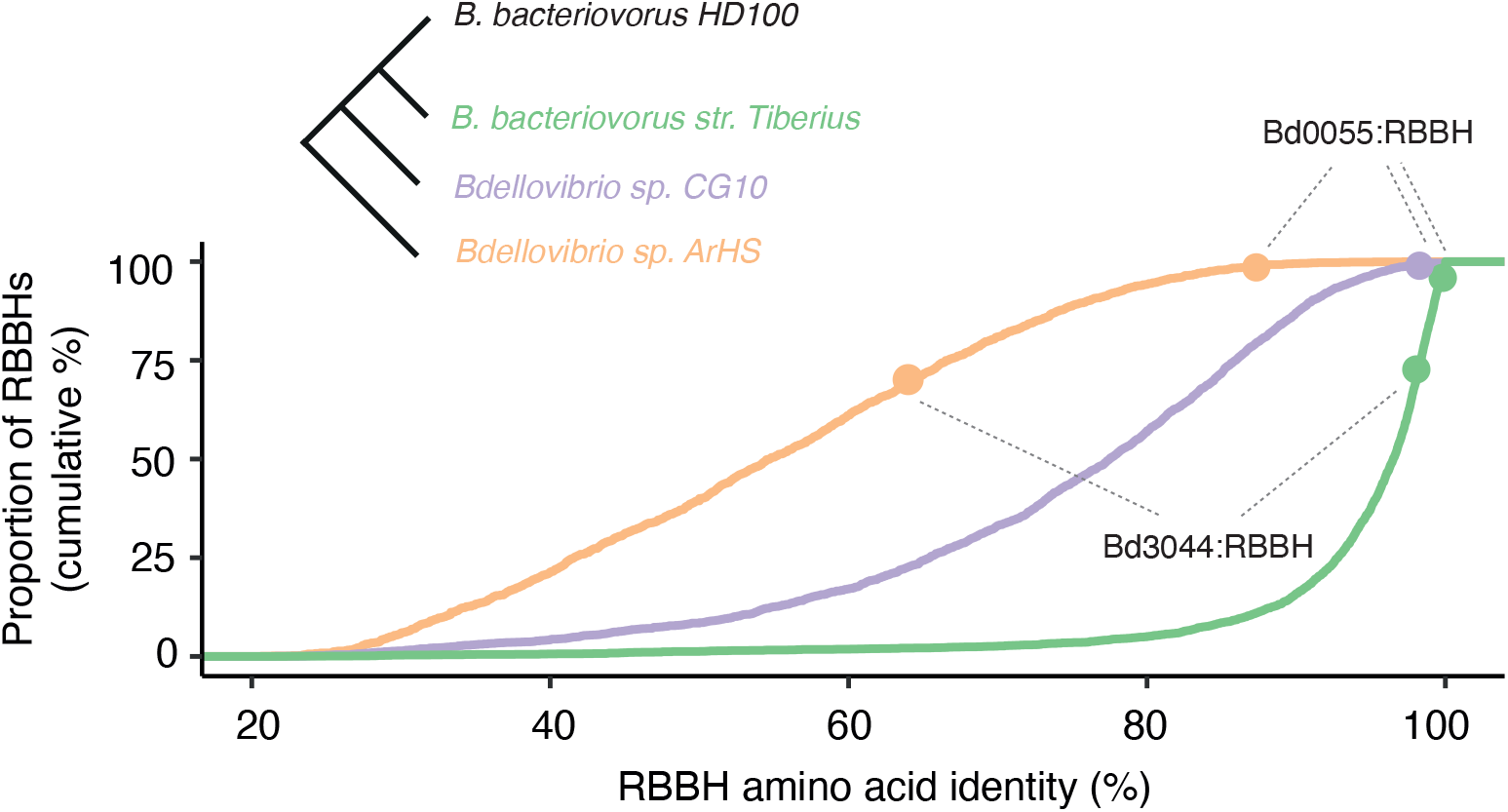
Conservation of Bd0055 at the amino acid level in the context of other proteins encoded by *B. bacteriovorus*. Homologs of proteins in *B. bacteriovorus* HD100 were identified in three other Bdellovibrio genomes of different divergence levels (*B. bacteriovorus* strain Tiberius, Bdellovibrio sp. CG10, and Bdellovibrio sp. ArHS) using a reciprocal best blast hit (RBBH) approach (see Methods). For each pairwise comparison, proteins were aligned and then ranked by amino acid identity. Bd0055 ranks amongst the most highly conserved proteins at different levels of proteome divergence. Bd3044 is comparatively less well conserved and absent from Bdellovibrio sp. CG10 (assembly ID: GCA_002773975.1).

**Figure S4.**
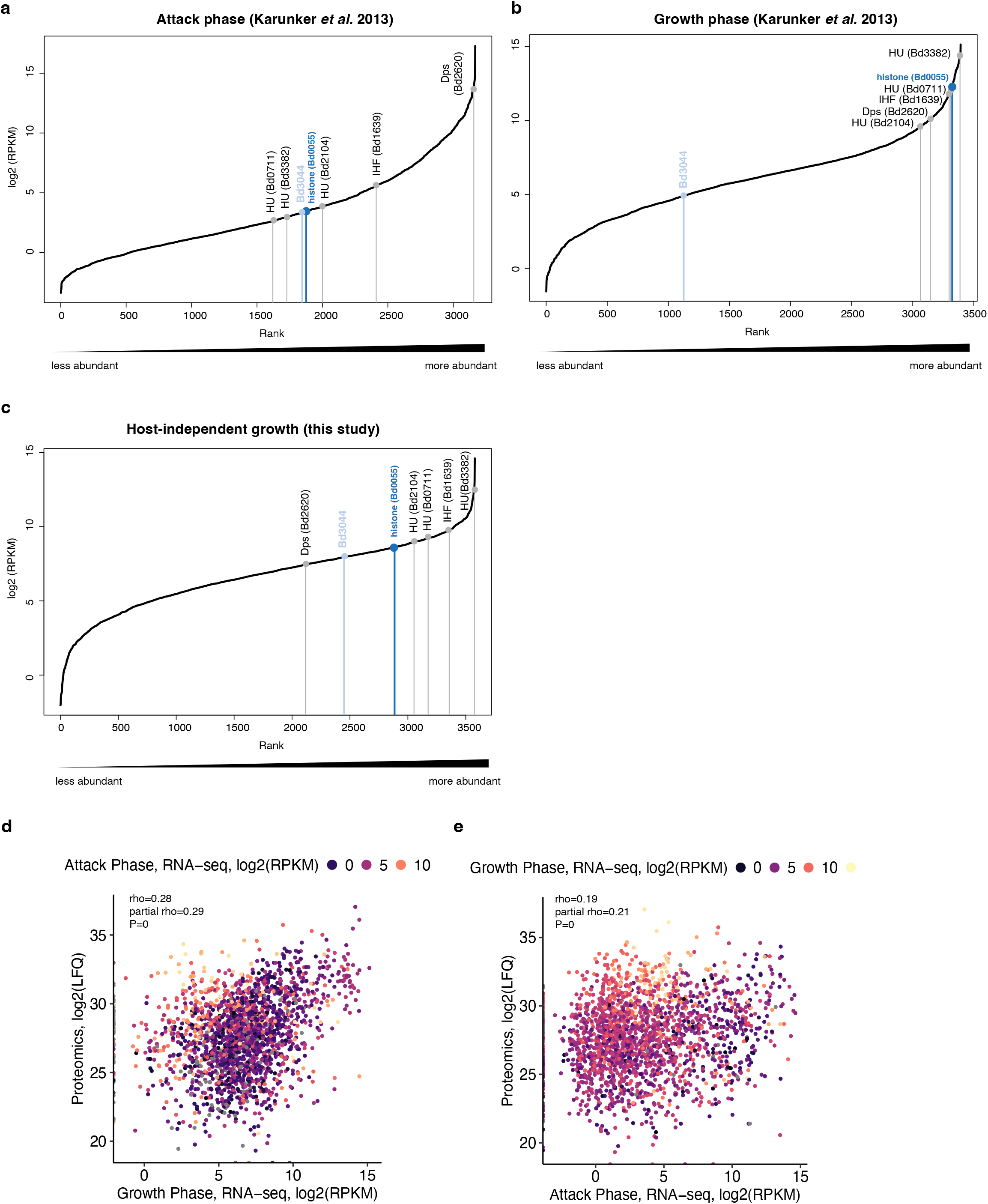
Ranked abundance of *B. bacteriovorus* transcripts during **(a)** attack phase, **(b)** growth inside the *E. coli* host, and **(c)** host-independent growth (strain HID13), highlighting transcript levels of Bd0055 relative to homologs of classic bacterial NAPs encoded in the **B. bacteriovorus** genome. **(d, e)** Correlation between protein abundance in attack phase cells (this study, whole cell extract) and RNA abundance during attack phase and growth inside the host (from Karunker *et al*. 2013).

**Figure S5.**
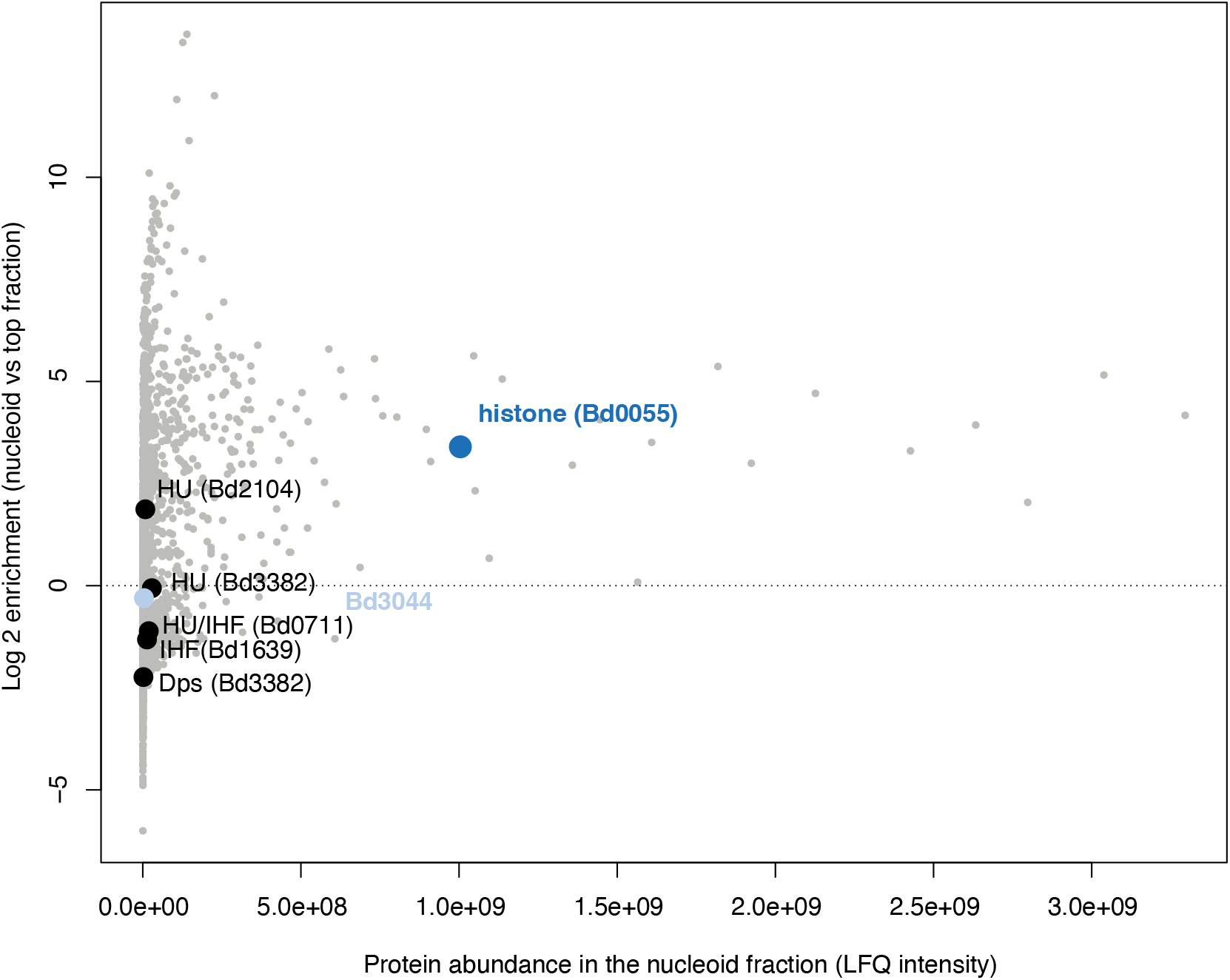
Protein abundance and nucleoid enrichment of Bd0055 in *B. bacteriovorus* in the context of ‘classic’ bacterial NAPs, for which homologs were detected in the *B. bacteriovorus* HD100 genome.

**Figure S6.**
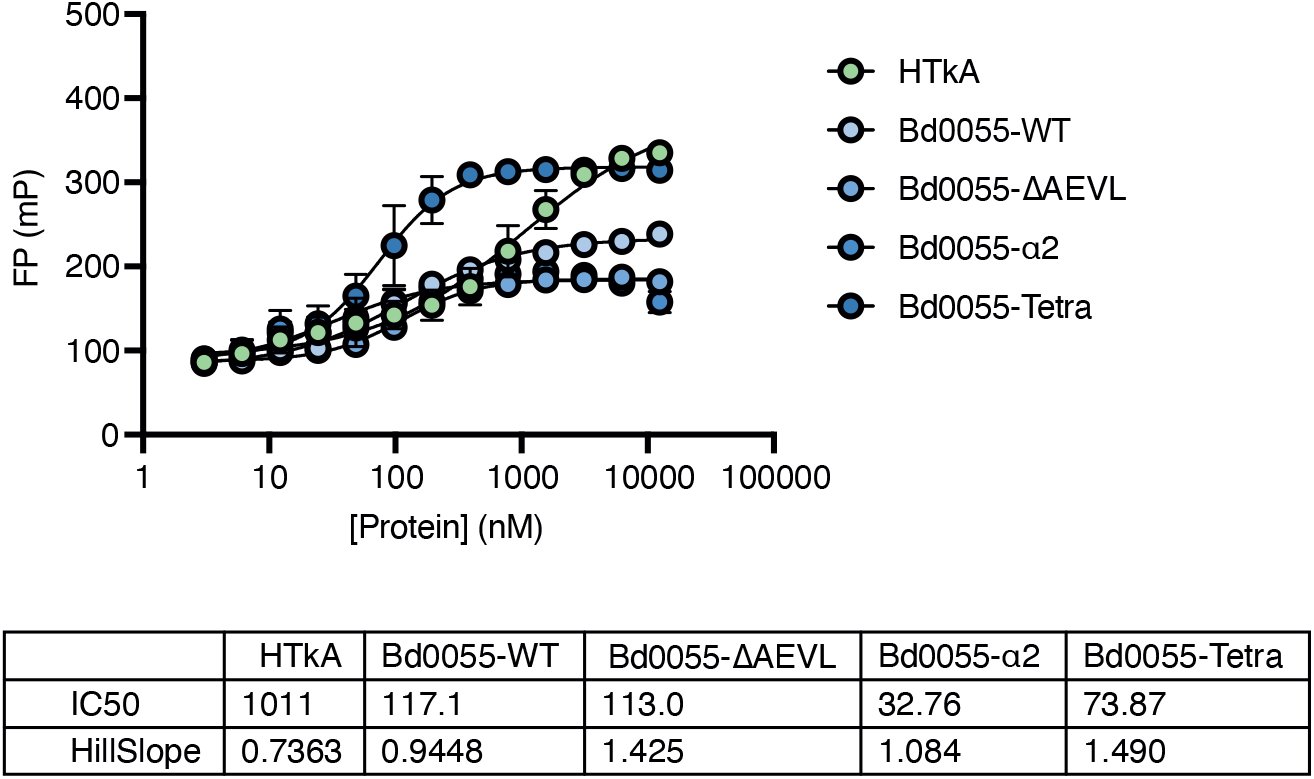
Fluorescence polarization (FP) of Bd0055 mutants binding to the same DNA as in Fig. 5d. FP is monitored using the Alexa488 flourophore (n=3). Error bars represent standard deviation of the mean.

**Figure S7.**
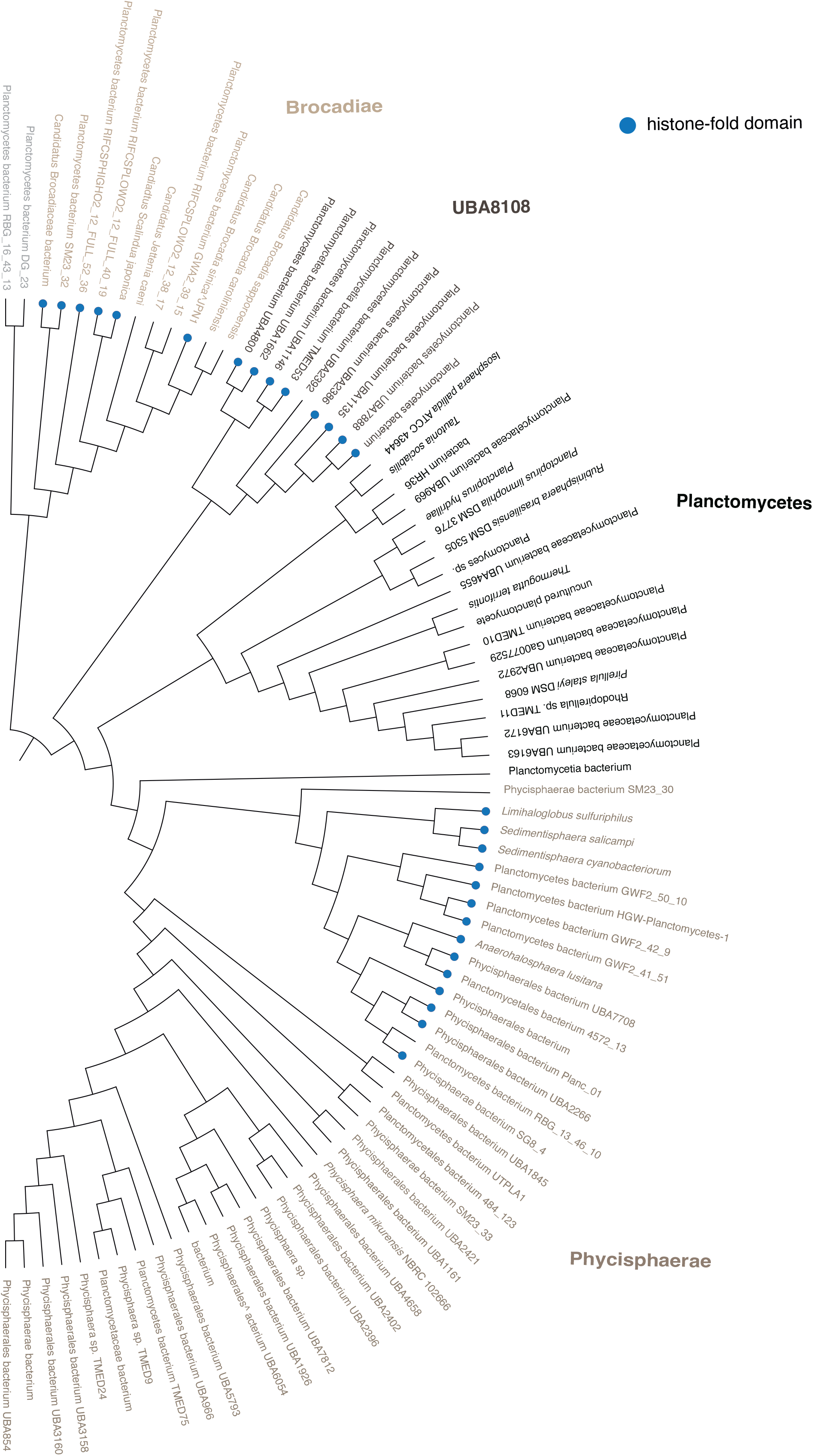
Cladogram of the Planctomycetota, indicating the presence of histone fold proteins throughout the clade. The cladogram is based on the publicly available GTDB tree (v 95.0) and represents the subset of genome assemblies that were used in this study and are also present in GTDB.

**Figure S8.**
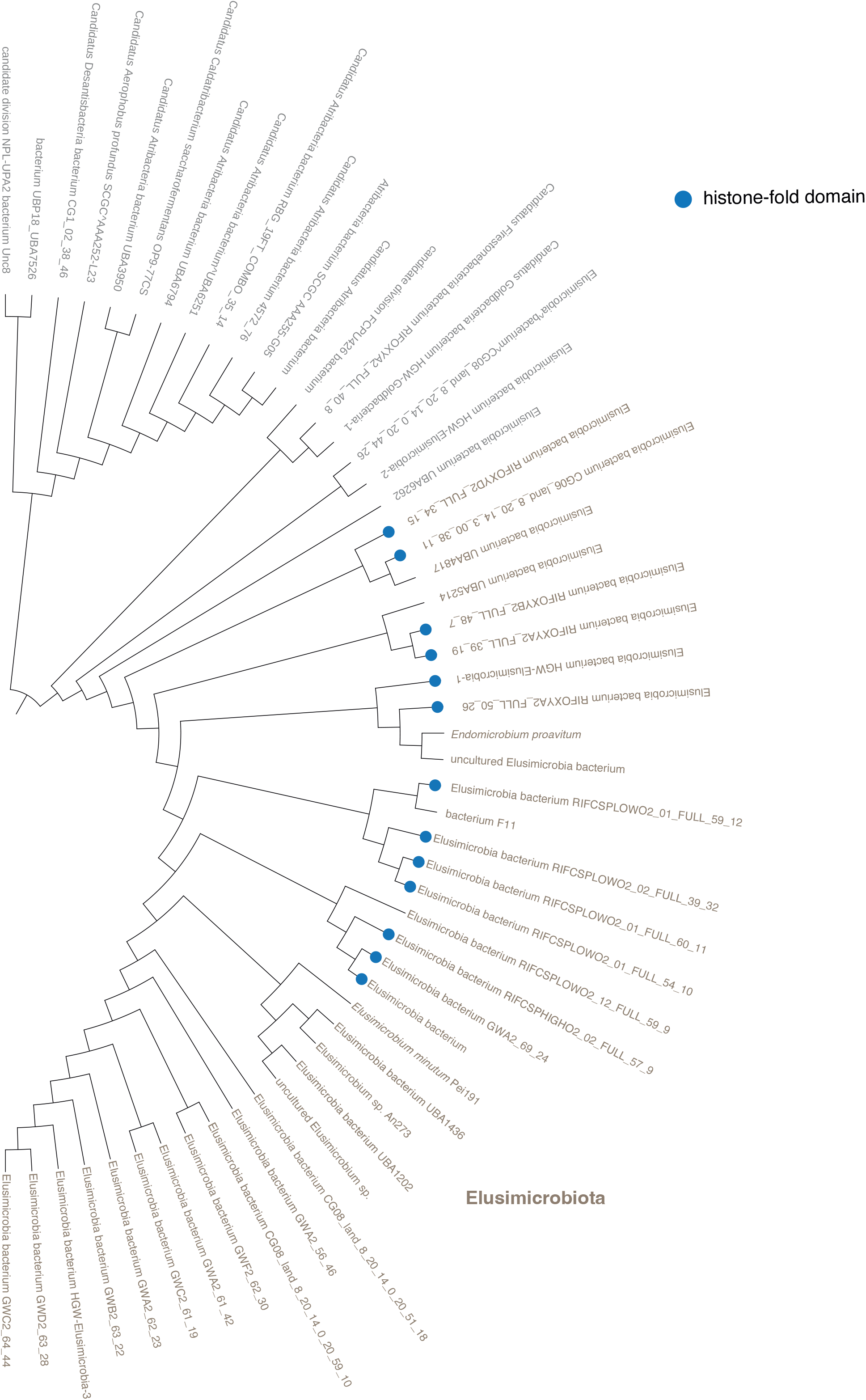
Cladogram of the Elusimicrobia, indicating the presence of histone fold proteins throughout the clade. The cladogram is based on the publicly available GTDB tree (v 95.0) and represents the subset of genome assemblies that were used in this study and are also present in GTDB.

**Figure S9.**
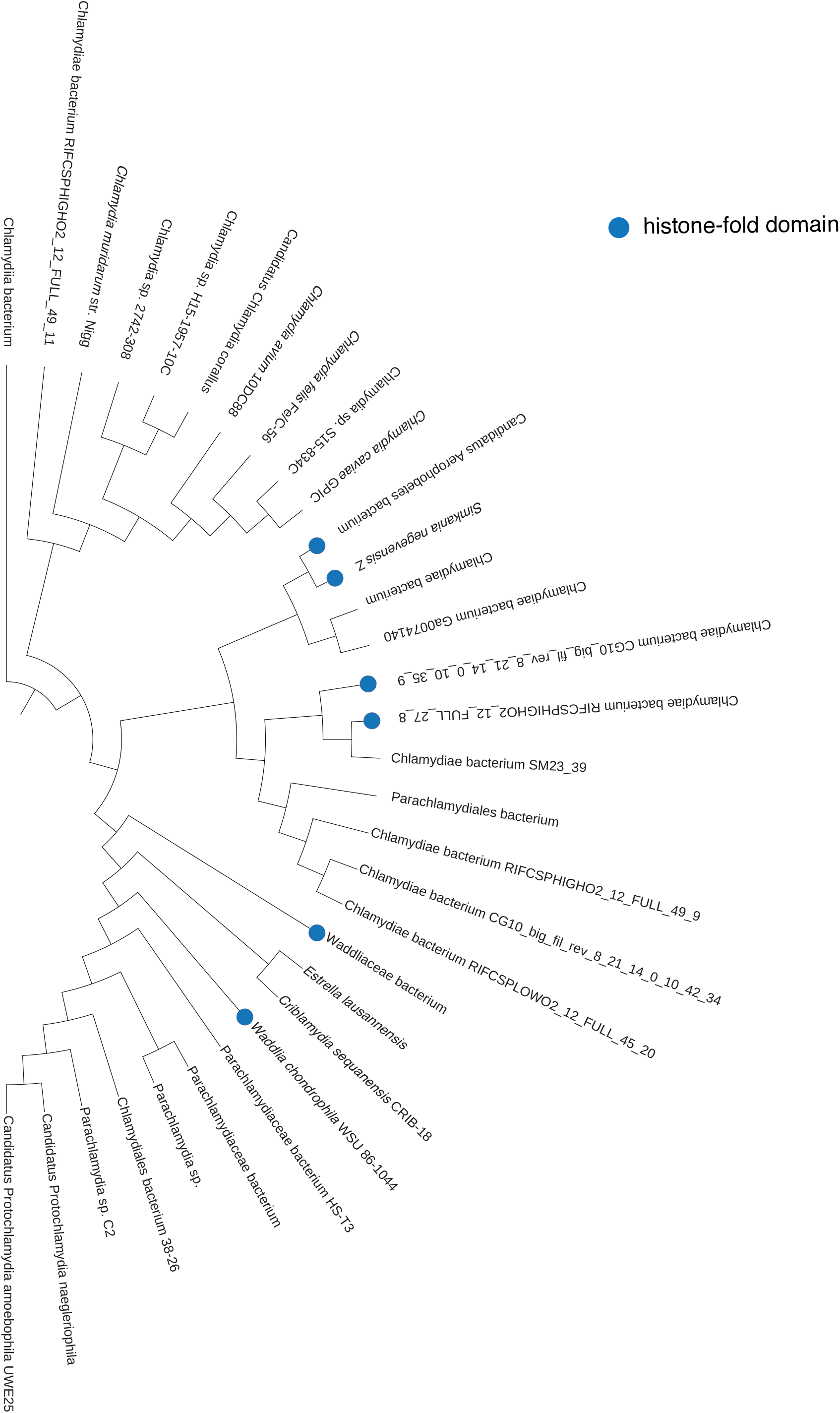
Cladogram of the Chlamydia, indicating the presence of histone fold proteins throughout the clade. The cladogram is based on the publicly available GTDB tree (v 95.0) and represents the subset of genome assemblies that were used in this study and are also present in GTDB.

**Figure S10.**
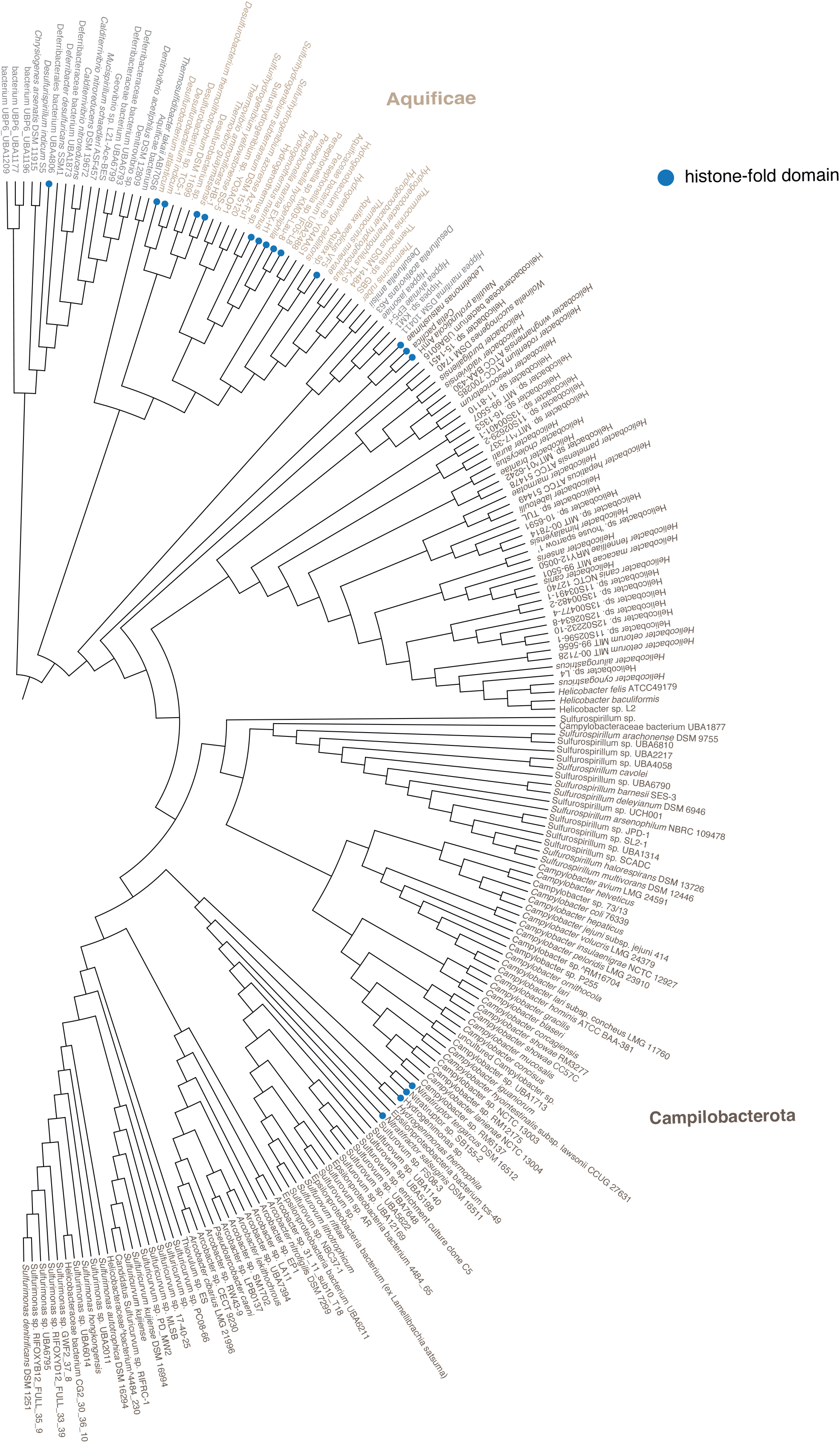
Cladogram of the Campilobacterota, Aquificae & sister clades, indicating the presence of histone fold proteins throughout the clade. The cladogram is based on the publicly available GTDB tree (v 95.0) and represents the subset of genome assemblies that were used in this study and are also present in GTDB.

**Figure S11.**
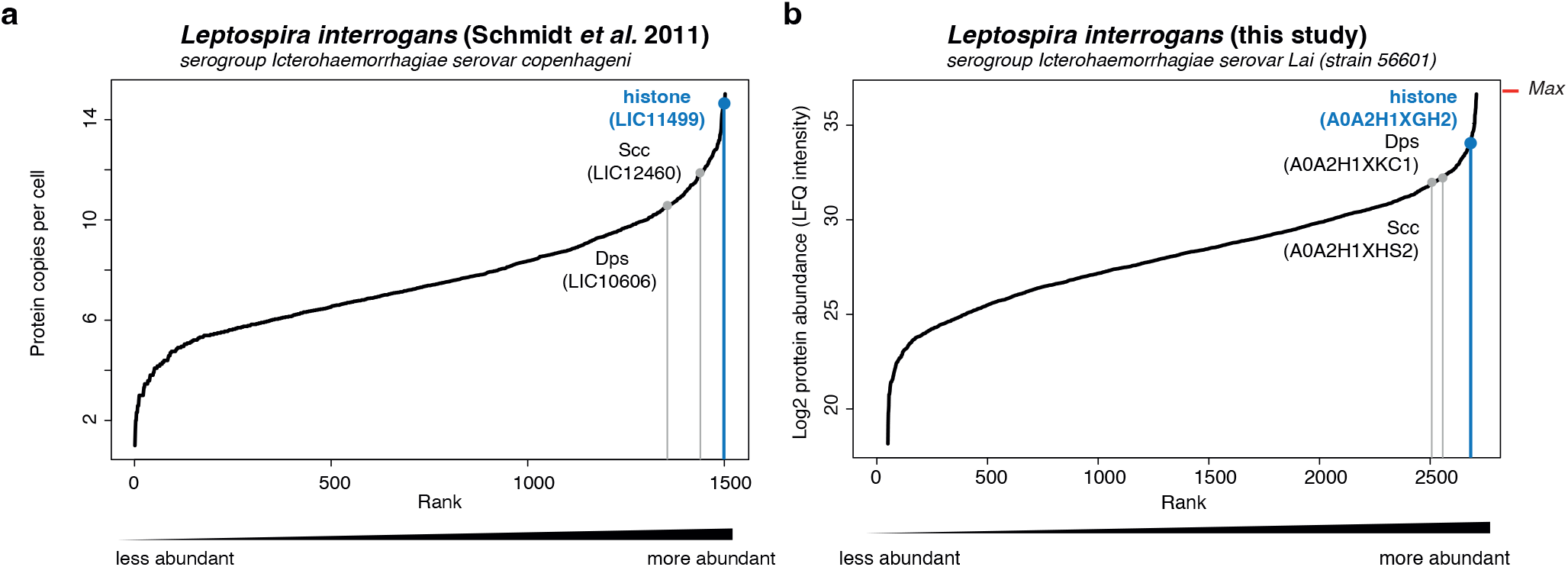
Ranked abundance of *L. interrogans* proteins from **(a)** Schmidt *et al*. (2011) and **(b)** this study, highlighting the abundance of the *L. interrogans* histone in the context of other known NAPs in this species. All quantified proteins are plotted.

**Figure S12.**
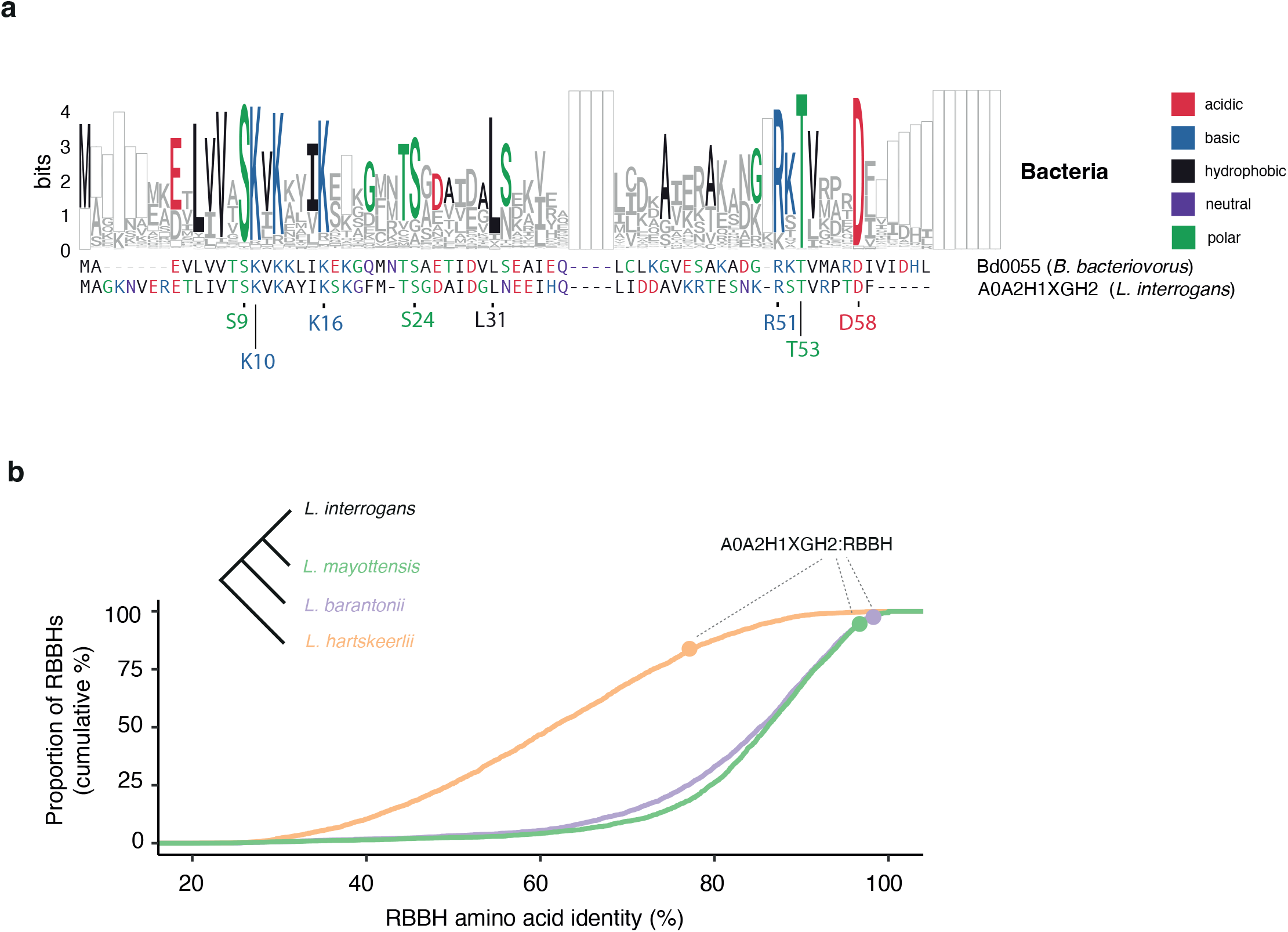
**(a)** Alignment of Bd0055 (*B. bacteriovorus*) and A0A2H1XGH2 (*L. interrogans*) in the context of overall amino acid usage across bacterial singlets (see Fig. 1c). **(b)** Conservation of the *L. interrogans* singlet histone (A0A2H1XGH2) at the amino acid level in the context of other proteins encoded by *L. interrogans*. Homologs of proteins in *L. interrogans* were identified in three other Leptospira genomes of different divergence levels (*L. mayottensis, L. barantonii, L. hartskeerlii*) using a reciprocal best blast hit approach (see Methods). For each pairwise comparison, proteins were aligned and then ranked by amino acid identity. A0A2H1XGH2 ranks amongst the most highly conserved proteins at different levels of proteome divergence.

**Figure S13.**
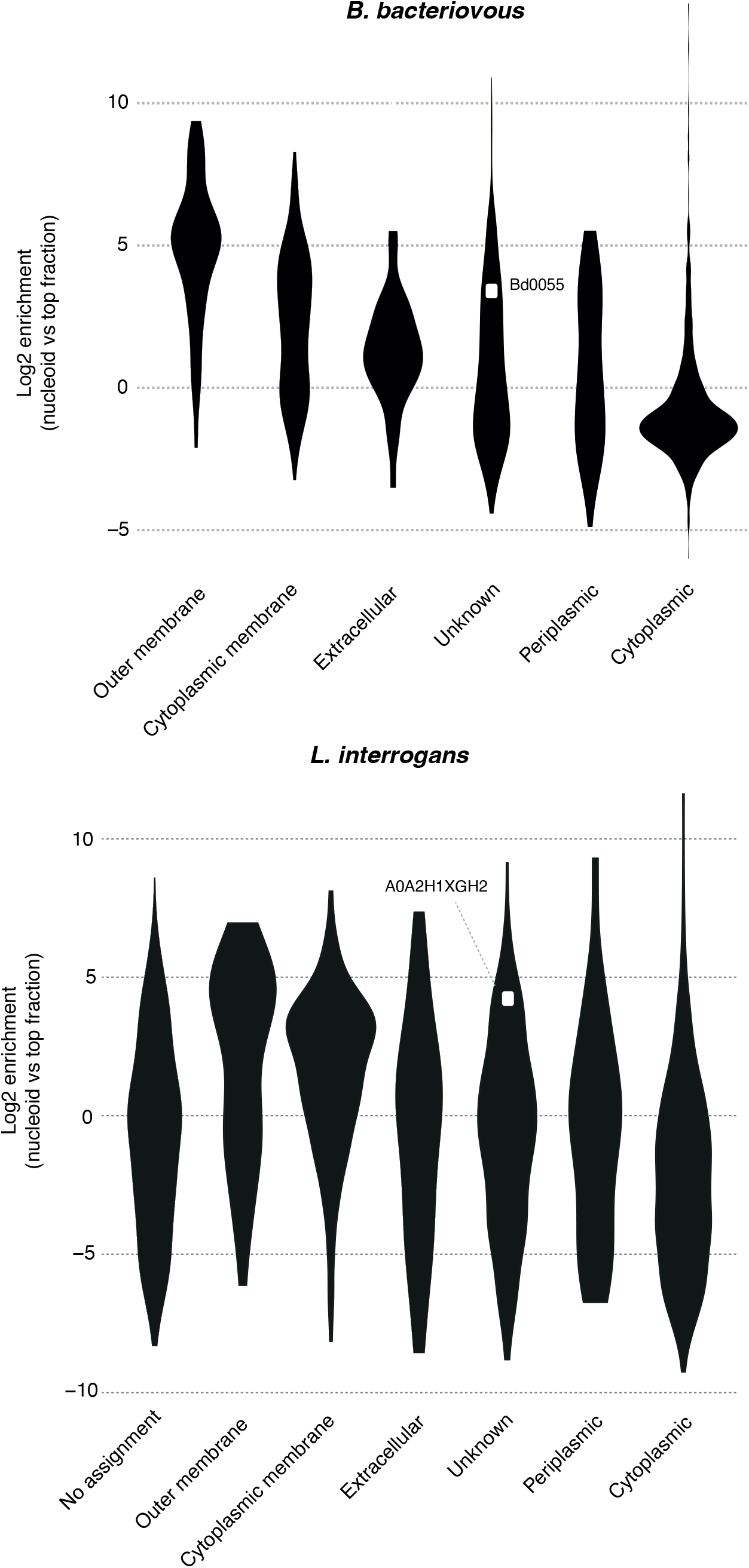
Nucleoid enrichment of proteins in *B. bacteriovorus* and *L. interrogans* as a function of their cellular localization as predicted by PSORTdb (https://db.psort.org/).

